# Pan-genomic and pan-transcriptomic analysis of the Heavy Metal ATPase family reveals diverse expression patterns and functional roles in barley

**DOI:** 10.64898/2026.07.07.736986

**Authors:** Jessica Shadbolt, Miriam Scheiber, Joanne Russell, Robbie Waugh, Kelly Houston

## Abstract

Heavy metals act as essential metalloprotein cofactors in numerous physiological processes but can become toxic when non-essential metals accumulate or when essential metals are in excess. As plants continuously encounter heavy metals through their roots, they have evolved complex homeostatic mechanisms to regulate metal uptake and distribution. The Heavy Metal ATPase (*HMA*) gene family encodes a group of heavy metal transporting P-type ATPases that have been linked to stress resistance and nutrient supply. Here, we used a bioinformatics approach to identify and characterise 13 *HMA* genes containing characteristic P_1B_-type ATPase domains and motifs in the barley Morex V3 reference genome. The genes are located on five of the seven barley chromosomes. Phylogenetic analysis revealed that they cluster into five sub-clades, including one clade unique to barley. Expression profiling across multiple datasets showed distinct temporal and tissue-specific expression patterns among *HvHMA*s, with several members exhibiting significant transcriptional responses to specific biotic and abiotic stresses. By utilising recently available pan-transcriptomic and pan-genomic resources, we have identified substantial allelic diversity and inter-accession variation in *HvHMA*s. Our findings suggest that *HvHMA*s have functions extending beyond canonical heavy metal homeostasis and warrant further investigation for their potential roles in broader physiological and stress-related processes.

## Introduction

Heavy metals are naturally occurring metallic elements characterized by their high atomic mass and density. Several of these elements, including copper (Cu), zinc (Zn), iron (Fe), manganese (Mn), nickel (Ni), and cobalt (Co), are classified as essential micronutrients required by all living organisms to sustain life [1]. An inadequate supply of these metal micronutrients can result in deficiency syndromes. These transition metals possess unique chemical properties that enable them to readily gain and lose electrons under physiological conditions. This allows them to participate in redox reactions and serve as critical cofactors in numerous enzymes involved in fundamental physiological processes such as respiration, photosynthesis, and nitrogen assimilation [2]. However, this same reactivity means that, at inappropriate concentrations, these heavy metals can promote the formation of reactive oxygen species, inducing oxidative stress and can displace native metals from their metalloprotein binding sites rendering them non-functional [3].

Increasing industrial activity, environmental pollution, and use of pig manure–based fertilisers (that contain high levels of Cu) have together contributed to the rising accumulation of heavy metals in the environment [4]. Plants, which are continually exposed to heavy metals through their contact with the soil, have evolved complex homeostasis mechanisms to ensure an appropriate supply of essential yet potentially toxic heavy metals [2]. Heavy Metal ATPases (*HMA*s), also known as P_1B_-type ATPases, are a conserved family of transmembrane transporters that enable plants to selectively traffic metal ions across cellular membranes by coupling transport with ATP hydrolysis (Li et al., 2015; Zhang et al., 2021). The first of these to be identified in plants and a canonical example of this transporter family is *At*HMA7 (*syn*. RESPONSIVE-TO-ANTAGONIST1 (*At*RAN1) [5] (**Fig. 1**). Like other P-type ATPases, P_1B_-type ATPases are multidomain proteins, consisting of a membrane (M) domain of multiple α-helices and several cytoplasmic domains. The cytoplasmic domains include an actuator (A) domain (containing the S/TGE motif), a phosphorylation (P) domain (containing the GDGxNDxP, and PxxK motifs), and a nucleotide binding (N) domain (**Fig. 1**) [6,7]. Structurally, P_1B_-type ATPases are distinguished from other P-type ATPases by the presence of eight transmembrane (TM) α-helices; the sixth of which contains a CPx (or SPC) motif proposed to be involved in cation translocation, and an HP locus in the N-domain associated with nucleotide co-ordination (**Fig. 1**). Typically, P_1B_-type ATPases also contain one or more putative metal binding domains at their N- or C-termini, which are referred to as Heavy Metal Associated (HMA) domains [8]. These domains, in addition to several structural features common to other P-type ATPases allow *HMA*s to transport heavy metals across cellular membranes. Like other P-type ATPases *HMA*s use an E1/E2 catalytic cycle to transport metal cations, whereby they transition between two main conformational states driven by ATP hydrolysis. Conformational changes enable the transport of bound metal ions through the transporter by alternately occluding access from either the cytosolic (or intravacuolar for organellar transporters) or extracellular side, allowing metal ions to be transported against concentration gradients [6,8].

**Figure 1:**
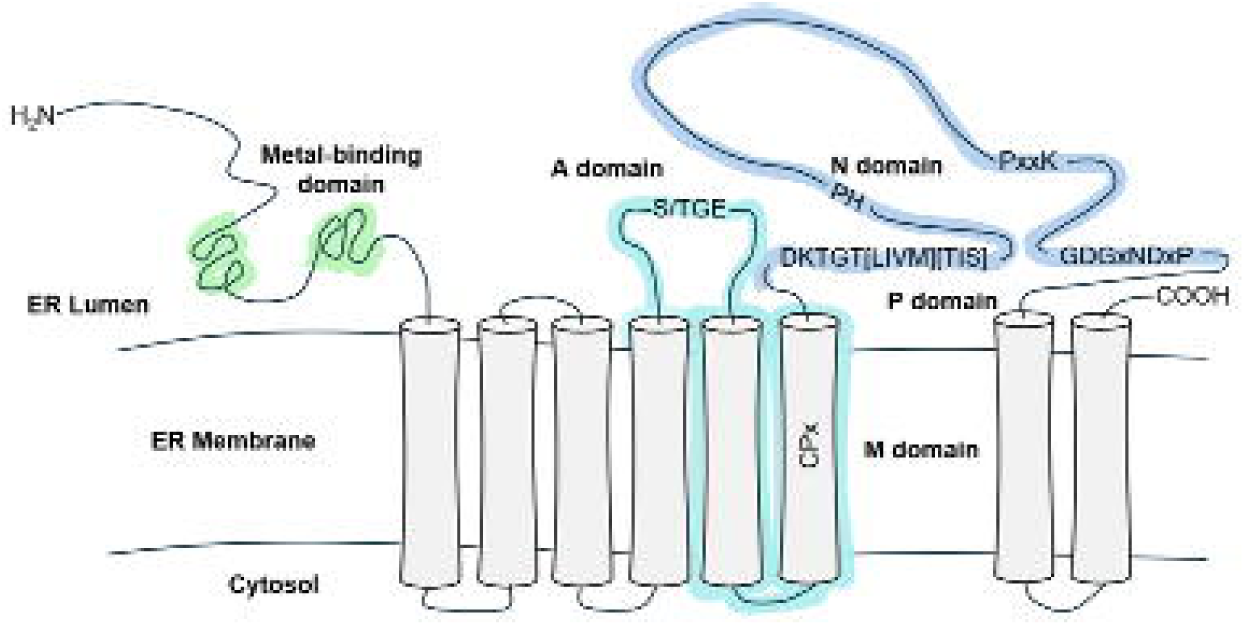
Schematic representation of the domain organisation and orientation of the canonical plant P_1B_-type ATPase *At*HMA7 (RAN1) located on the Endoplasmic Reticulum (ER) membrane. Regions corresponding to the HMA, E1–E2 ATPase, and hydrolase domains are highlighted in green, cyan, and blue, respectively. Cylinders indicate the positions of the eight transmembrane helices. Conserved sequence motifs are shown as letter strings (S/TGE, CPx, DKTGT[LIVM][TIS], HP, PxxK, and GDGxNDxP) located within the membrane (M), actuator (A), phosphorylation (P), and nucleotide-binding (N) domains.

Extensive study of *HMA* family members is warranted by their diverse roles in micronutrient nutrition, detoxification, and phytoremediation [9–11]. In crops, *HMA*s are particularly significant as they have been linked to metal accumulation in edible tissues [12–14], metal stress tolerance, and drought tolerance [15]. Additionally, there is increasing evidence linking *HMA*s with plant immunity [16]. Given their physiological significance and potential economic impact, the *HMA* gene family has been studied across multiple crop species and their progenitors, including wheat (*Triticum aestivum*) [17], goatgrass (*Aegilops tauschii*) [18], tomato (*Solanum lycopersicum*) [15], rice (*Oryza sativa*), maize (*Zea mays*), sorghum (*Sorghum bicolour*) [19], rapeseed (*Brassica napus*) [20], and soybean (*Glycine max*) [21]. Increased genomic resources in plants have broadened the characterization of the *HMA* gene family and, together with higher-quality genome assemblies, have enabled the discovery (or removal) of additional family members.

Although several *HMA* genes have been characterized through mutant studies and heterologous expression assays in other species, barley *HMA* genes (*HvHMA*s) have yet to receive comparable attention. This is surprising given that, among cereal crops, barley is particularly resilient and can be cultivated on nutrient-poor and marginal soils where other cereals cannot thrive, making its regulation of essential metal micronutrients especially intriguing [22]. The first detailed identification of a member of the *HMA* gene family in barley was by Mills *et al.* (2012), who reported nine *HvHMA* genes and characterised *HvHMA2*. This was later expanded by Zhang *et al.* (2021) who identified 21 *HvHMA* family members and performed expression analyses on five of these genes in the leaves of seedlings under Cd stress [23].

Since then, new genomic and transcriptomic resources have become available for barley, providing the opportunity for more comprehensive analyses. In this study we interrogated the most up-to-date barley reference genome assembly to identify members of the *HMA* gene family [24]. We then explored pan-genomic and pan-transcriptomic resources to assess sequence variation and expression diversity among barley accessions, revealing intraspecific differences in both sequence and expression [25,26]. Finally, we analysed *HvHMA* expression patterns across developmental stages, in a range of tissues, and under multiple biotic and abiotic stress conditions using publicly available expression datasets [25].

The barley pan-genome includes 17 cultivars, 36 landraces and 23 wild barleys whereas the barley pan-transcriptome contains a subsample of eight cultivars, 11 landraces and one wild barley. These overlapping resources therefore capture both barley genetic and functional diversity. As a result, our findings expand our understanding of the *HMA* gene family in barley and highlight the extensive natural variation present within this species, including alleles that may hold promise for optimising metal homeostasis and contribute to future breeding programs.

## Results

### Genome-wide identification and classification of barley HMAs

We identified 13 *Hv*HMA sequences within the Morex v3 barley genome sequence assembly [24] (**Supplementary Table S1**). To assign *Hv*HMA names, the 13 *Hv*HMA sequences were aligned with known HMA proteins from Arabidopsis, wheat, and rice (**Supplementary Table S2**). Overall, the *Hv*HMAs showed the highest sequence similarity to wheat orthologues, with identity ranging from 15.7% to 98.9% (**Supplementary Table S2**). As a result, *Hv*HMA names were assigned primarily based on protein sequence similarity to named wheat HMAs. Thus, for seven of the identified *Hv*HMA sequences clear orthologues in wheat could be identified (> 92% sequence identity), and these sequences were assigned *Hv*HMA1, 3–6, 8, and 9 (**Supplementary Table S2**). Two barley sequences, HORVU.MOREX.r3.7HG0727750 and HORVU.MOREX.r3.7HG0727780, displayed relatively high conservation to wheat orthologues *Ta*HMA2;5 and *Ta*HMA2;2 (94.2% and 91.1%, respectively, **Supplementary Table S2**). The physical proximity of these sequences on chromosome 7H (**Fig. 2a**) (121,883 bp between the end of the final exon of HORVU.MOREX.r3.7HG0727750 and the start of the first exon of HORVU.MOREX.r3.7HG0727780), coupled with their high sequence similarity to one another relative to other *Hv*HMAs (68.9%, **Supplementary Table S3**) suggests they likely arose from a gene duplication event and were therefore designated as *Hv*HMA2a and *Hv*HMA2b.

**Figure 2:**
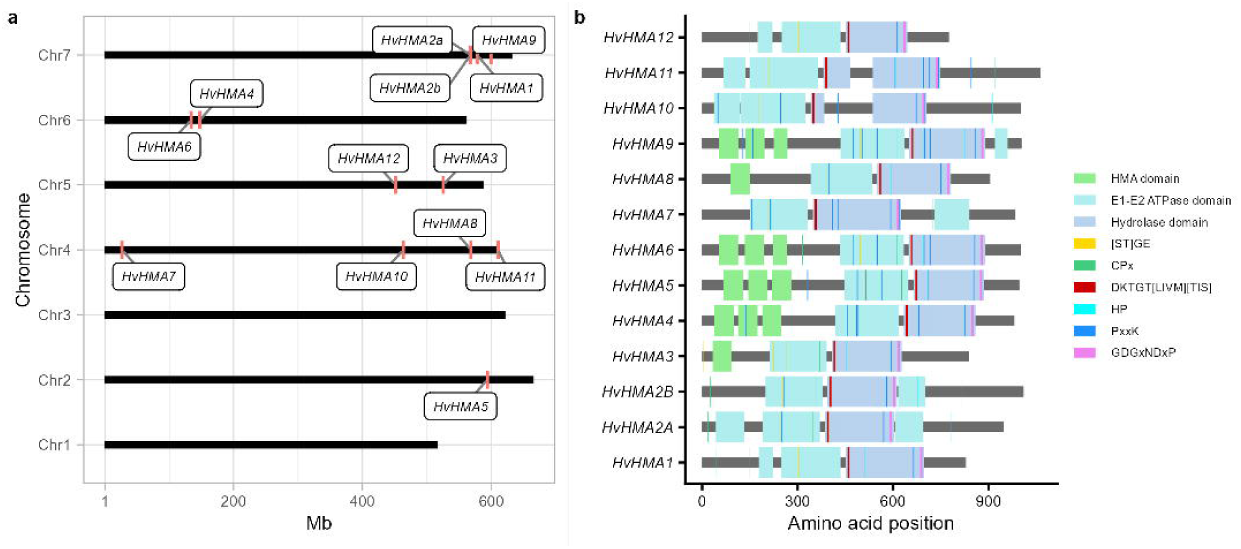
Members of the *HMA* gene family in barley. (A) Chromosomal positions of *HMA* family members in barley. The X-axis represents the physical locations of the *HvHMA* genes, shown in red, on chromosomes according to the Morex v3 physical map (Mascher, 2021). (B) Schematic representing the positions of the functional domains and defining motifs present within the barley *HvHMA* gene family. The colour coding corresponds to the domains and motifs key shown within the diagram.

The remaining *Hv*HMA sequences, HORVU.MOREX.r3.4HG0419450, HORVU.MOREX.r3.4HG0339640, and HORVU.MOREX.r3.4HG0386480, exhibited relatively low conservation with HMA sequences from any of the other analysed species (ranging from 3.6% to 18.8% identity, **Supplementary Table 2**). HORVU.MOREX.r3.4HG0419450 was designated *HvHMA7*, as it showed the highest sequence identity (15.1%) to *Ta*HMA7. HORVU.MOREX.r3.4HG0339640 and HORVU.MOREX.r3.4HG0386480 were assigned *Hv*HMA10 and *Hv*HMA11 based arbitrarily on the 5’ to 3’ position of their associated gene models on chromosome 4H. Finally, HORVU.MOREX.r3.5HG0484850 was designated *Hv*HMA12. In this study, naming of *Hv*HMAs based on orthology with wheat HMAs provided some consistency with previous conventions—particularly those reported by Mills *et al.* (2012). However, discrepancies remain, and the *Hv*HMA names used here may not correspond directly to those previously reported for the same sequences (**Supplementary Table 1**) [23].

### Molecular characteristics of *HMA* proteins and chromosomal location of *HMA* genes in barley

*HvHMA* gene length ranged from 3,097 bps (*HvHMA3*) to 34,617 bps (*HvHMA12*) and their CDS ranged from 2,560 to 3,929 bps (**Supplementary Table 1**). All *HvHMA* genes contained at least six exons, with the greatest number of exons (34) in *HvHMA7*. Barley *HMA* genes were located on chromosomes 2H and 4-7H (**Fig. 2a**). Protein lengths of *Hv*HMA1-13 ranged from 774 (*Hv*HMA12) to 1,062 (*Hv*HMA10) amino acids and weighed between 82.02 kDa (*Hv*HMA12) and 115.87 kDa (*Hv*HMA10). The theoretic isoelectric points of the barley *HMA* family were mostly acidic, from 4.99 (*Hv*HMA6) to 7.94 (*Hv*HMA12). All *Hv*HMA proteins were predicted to localise to the plasma membrane, except *Hv*HMA3 which was predicted to localise to the vacuole (**Supplementary Table 1**).

### Domain annotation and motif distribution within barley HMA proteins

To be defined as *HMA* family member, all proteins needed to contain at least one occurrence of the following motifs: S/TGE, CPx (or SPC), DKTGT[LIVM][TIS], HP, PxxK, and GDGxNDxP. The positions of these motifs within the identified *Hv*HMA proteins are indicated in **Fig. 2b**. Motifs PxxK, CPx (or SPC), and HP occurred in many of the *Hv*HMA proteins at multiple positions. The *Hv*HMA proteins identified in this study are composed of at least one E1-E2 ATP-ase domain followed by a Hydrolase domain. In all the barley *HMA* proteins, the P- and A- domain conserved regions DKTGT[LIVM][TIS], TGD, PxxK, and GDGxNDxP [8,27] were present in the same order within the identified hydrolase domain, outlining the putative active site region for Mg^2+^ and phosphate binding [27]. Additionally, *Hv*HMA3-6, *Hv*HMA8, and *Hv*HMA9 contained one to three copies of the Heavy Metal Associated (HMA) domain within the first 300 residues of the N-terminus of the protein.

### Structural prediction of the *Hv*HMA family in Morex

To gain insight into the three-dimensional architecture of the *Hv*HMA family, protein structures were predicted for these genes in the reference cultivar Morex and visualised with residues coloured according to the confidence of the structural predictions (**Fig. 3**). Broadly, the structure of the *Hv*HMAs was consistent, with a relatively more conserved core of eight α-helices forming the canonical membrane spanning pore, and more diverse cytosolic domain which contained both helices and β-pleated sheets (**Fig. 3**). Supporting their predicted membrane localisation, all *Hv*HMA structures exhibited a distinct membrane-spanning region corresponding to the transmembrane domain. In *Hv*HMA3 and *Hv*HMA12 (**Fig. 3d** and **3m**, respectively), the pore structure was less conserved, and the predicted models for these family members showed putative transmembrane helices that appeared truncated or oriented inconsistently with the canonical pore architecture. Multiple *Hv*HMA proteins, including *Hv*HMA2a, *Hv*HMA2b, and *Hv*HMA3 (**Fig. 3b-d**), exhibited long regions with relatively low confidence at their C-terminus which appear disordered due to their lack of conservation with other known structures. These C terminal regions are all predicted to be oriented on the cytosolic side of the transporter and their sequence-level divergence may be indicative of neofunctionalization of this gene family in barley.

**Figure 3:**
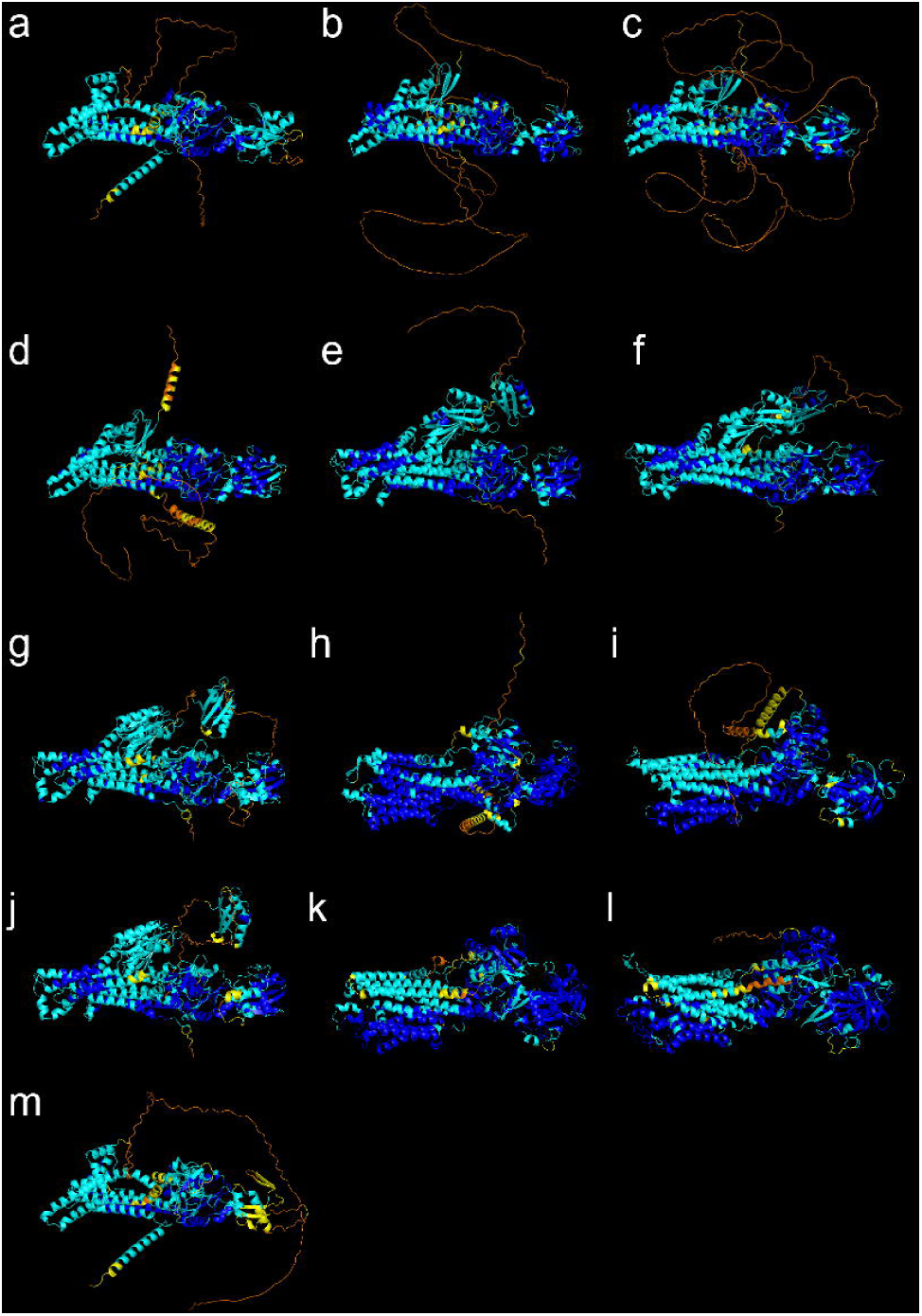
Predicted structures of the Morex HvHMA proteins. Residues are coloured by their predicted local distance difference test (pLDDT) score, with very high confidence (pLDDT > 90) coloured in blue, confident (70 < pLDDT ≤ 90) in cyan, low confidence (50 < pLDDT ≤ 70) in yellow, and very low confidence (pLDDT ≤ 50) in orange. (a) *Hv*HMA1, (b) *Hv*HMA2a, (c) *Hv*HMA2b, (d) *Hv*HMA3, (e) *Hv*HMA4, (f) *Hv*HMA5, (g) *Hv*HMA6, (h) *Hv*HMA7, (i) *Hv*HMA8, (j) *Hv*HMA9, (k) *Hv*HMA10, (l) *Hv*HMA11, and (m) *Hv*HMA12.

### Phylogenetic analysis of the *HMA* family across multiple species

To explore the evolutionary relationships between barley, wheat, Arabidopsis, sorghum, maize, and rice HMAs, protein sequences for 13 *HvHMA*s, 29 *TaHMA*s, nine *OsHMA*s, 11 *SbHMA*s, 11 *ZmHMA*s, and six *AtHMA*s were used to construct a phylogenetic tree (**Supplementary Fig. S1**). This indicated that the *HMA*s were distributed into five subgroups (A-E) with *Sb*HMA7 forming an outgroup. Within the subgroups, four *Hv*HMAs belong to group A, one to group B, three to group C, two to group D, and three to group E. In most cases HMA orthologues from different species clustered within the same subgroup: clade A contained all HMA4, HMA5, HMA6 and HMA9 members, clade B contained all HMA8 members, clade C contained all HMA2 and HMA3 members, and clade D contained all HMA1 members (**Supplementary Fig. S1**). Monocotyledonous species (rice, wheat, sorghum, and maize) were more closely related than *HMA* members from dicotyledonous Arabidopsis (**Supplementary Fig. S1**). In contrast, HMA7 orthologues are spread across clades A, B, and E due to the relatively low sequence conservation between HMA7 sequences from these clades (**Supplementary Table S2**). Clade E of the phylogenetic tree consists of only barley HMAs (*Hv*HMA7, *Hv*HMA10, and *Hv*HMA11). The position of these HMAs on the phylogenetic tree is due to their diverse sequences in comparison to other family members and indicates that these members do not have close homologs amongst the analysed HMA proteins.

### *HMA* allele diversity across the Barley Pangenome v2

The recent release of whole genome sequencing data across 76 diverse barley accessions as part of the Barley Pangenome v2 project (BPv2) has allowed us to assess the diversity of the *HvHMA* family at the species level. A table summarising the origin, growth habit, row-type, and status of the BPv2 genotypes is provided in **Supplementary Table S4**. **Figure 4** presents bar plots showing the number of alleles identified for each *HvHMA* gene and the proportion of accessions carrying each allele, while **Supplementary Tables S5–S17** detail the allele present in each surveyed accession. Homologues of each *HMA* gene family member were identified in almost all the 76 BPv2 genotypes. Homologues of *HvHMA2b* and *HvHMA11* could not be identified in accessions WBDC133 and HOR495, respectively. Furthermore, homologues of *HvHMA12* could not be identified in 27 of the analysed accessions. Accessions in which *HMA* homologues could not be identified were predominantly wild or landrace types, which we observed were more likely to carry alleles with diverse sequences. Therefore, it is possible that the alleles present are insufficiently conserved to be detected under the thresholds used in our BLAST search. The allelic diversity of *HvHMA* genes varied widely, with the number of identified alleles ranging from 3 (*HvHMA10*) to 32 (*HvHMA2a*) (**Supplementary Tables S15** and **S6**). While most genes were dominated by a single allele, *HvHMA2a*, *HvHMA3*, and *HvHMA9* exhibited increased allelic diversity. Except for *HvHMA4*, the number of alleles for each *HMA* gene increased across the groups—cultivars, landraces, and *H. spontaneum* (wild barley) (**Fig. 4**). The number of *HvHMA4* alleles was higher in cultivars (five alleles across a total of 17 cultivars) than in landraces (two alleles across a total of 36 landraces) (**Fig. 4**, **Supplementary Table S9**). All *HvHMA* genes have a single copy in this dataset.

**Figure 4.**
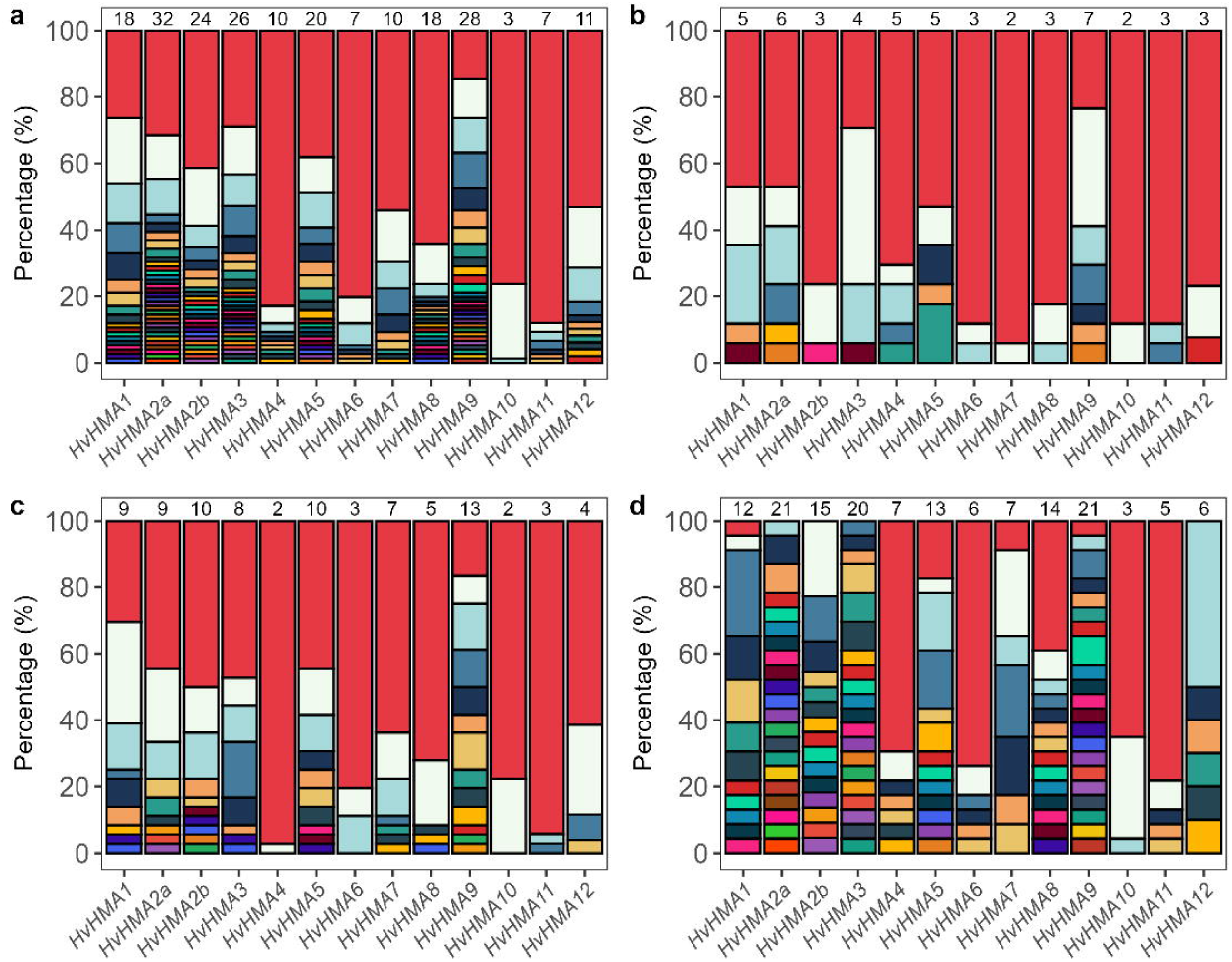
Allele diversity of *HvHMA* genes across 76 barley pan-genome (BPv2) genotypes. Stacked bar plots show the relative frequency (percentage of total identified homologues) of unique *HvHMA* alleles based on predicted protein sequences. Each colour represents a distinct allele, with red indicating the most frequent (major) allele for each gene. Numbers above each bar denote the total number of unique alleles identified. Panels represent (a) all 76 BPv2 genotypes, (b) 17 cultivated accessions, (c) 36 landraces, and (d) 23 *Hordeum spontaneum* (wild) accessions.

### Several barley genotypes carry frameshift *HMA* alleles, including elite cultivar Barke

Within the pangenome collection, alleles containing frameshift-inducing indels relative to the Morex sequence were identified in *HvHMA1*, *HvHMA2a*, *HvHMA2b*, *HvHMA4*, *HvHMA7*, *HvHMA8*, and *HvHMA11* (**Supplementary Table S18**). The impact of these polymorphisms depends on their position within the coding sequence and whether the reading frame is restored by compensatory indels [28,29]. In *Hv*HMA1, genotypes B1K-33-13, HOR2180, and HOR12184 contain frameshifts beginning at amino acid positions 21, 42, and 53, respectively, resulting in premature stop codons and proteins lacking both the hydrolase and E1–E2 ATPase domains (**Supplementary Fig. S2a**). A frameshift allele of *Hv*HMA2a was identified in OUN333. Multiple insertions and deletions transiently restore the reading frame at amino acid position 202, but a subsequent premature stop codon truncates the predicted protein at amino acid 214 within exon 4. This restored region corresponds to only a small portion of the E1–E2 ATPase domain (**Supplementary Fig. S2b**). No other HMA frameshift allele was identified with reading-frame restoration through similar compensatory indels. Frameshift alleles of *Hv*HMA2b were identified in HOR18321 and HOR21256. In HOR18321, a frameshift at position 25 results in truncation at amino acid 67, whereas in HOR21256 a frameshift at position 381 causes truncation at position 388, leaving most of the preceding E1–E2 ATPase domain intact (**Supplementary Fig. S2c**). Alleles with indels causing frameshifts were identified in *Hv*HMA7, *Hv*HMA8, and *Hv*HMA11. HOR12184 carries an *Hv*HMA7 allele truncated at amino acid 162 following a frameshift at position 103, HOR7172 carries an *Hv*HMA8 allele truncated at position 81 following a frameshift at position 24, and HOR12541 carries a *Hv*HMA11 allele truncated at position 203 following a frameshift at position 162 (**Supplementary Fig. S2d-f**). Among the HMA genes, *Hv*HMA4 had the greatest number of frameshift alleles (**Fig. 5a**), with four distinct variants identified from the sequencing data. Genotypes B1K-17-07, WBDC133, HOR4224, and HOR14061 share the same frameshift allele, whereas Barke, HID84, and HOR2180 each carry unique variants. The HID84 allele is predicted to be the most severely affected, with a frameshift at amino acid position 546 resulting in complete loss of the hydrolase domain (**Fig. 5a**). In contrast, the remaining three frameshift alleles, including that present in the elite cultivar Barke, are truncated within the GDGxNDxP motif, with the frameshift occurring at the variable "x" residue (**Fig. 5a**). Evidence of *in planta* expression data (the analysis of which is described in following sections) was available for only two frameshift alleles: *HvHMA2a* in OUN333 and *HvHMA4* in Barke. The presence and expression of a frameshift allele in the widely cultivated elite variety Barke, together with the functional importance of *Hv*HMA4, motivated further structural analysis of the GDGxNDxP motif.

**Figure 5.**
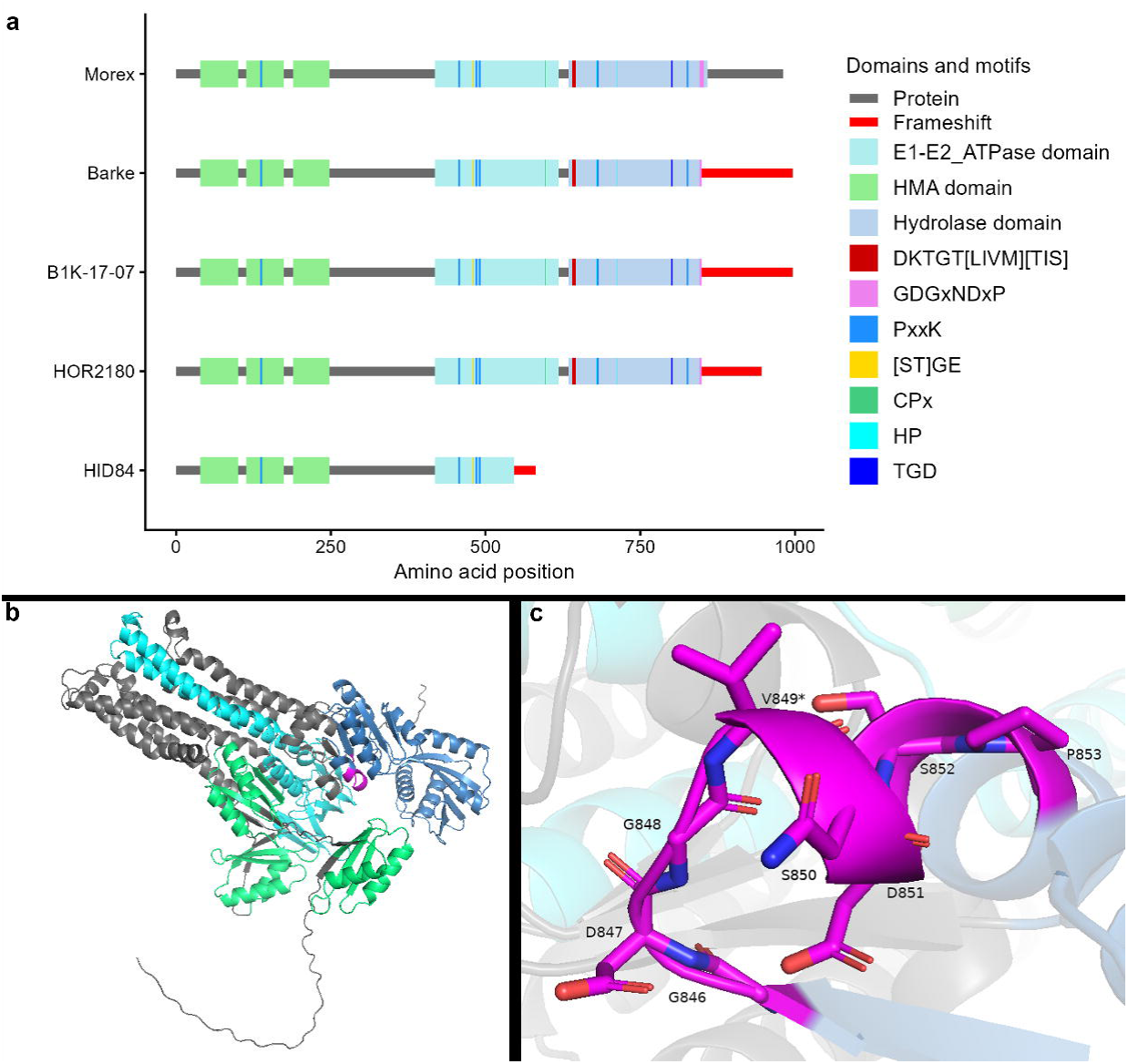
Multiple frameshift alleles of *HvHMA4* are present in barley pangenome accessions. (a) Schematic comparison of the predicted protein products encoded by *HvHMA4* alleles from selected barley accessions and the reference cultivar Morex. Grey bars represent the protein sequence, coloured boxes indicate conserved domains and motifs, and red regions denote amino acid sequences encoded downstream of the frameshift mutation. The positions of the HMA domain, E1–E2 ATPase domain, hydrolase domain, and conserved HMA motifs are shown according to the legend. Frameshift alleles result in truncation of the C-terminal region and loss of conserved motifs to varying extents. Amino acid positions are indicated on the x-axis.(b) Predicted structure of the Morex *Hv*HMA4 protein. Conserved domains are coloured as in panel A, with the GDGxNDxP motif highlighted in magenta. (c) Close-up view of the GDGxNDxP motif (also known as the hinge motif) within the predicted *Hv*HMA4 structure. Conserved residues forming the motif are shown as sticks and labelled. Oxygen atoms are labelled here in red and nitrogen atoms in blue. The frameshift mutations identified in the pangenome accessions B1K-17-07, WBDC133, HOR4224, HOR14061, Barke, and HOR2180 occur at the valine residue (V849) demarked with an asterisks (*), disrupting the motif and resulting in loss of the downstream conserved C-terminal domain.

The position of the GDGxNDxP motif in Morex *Hv*HMA4 relative to the HMA, E1-E2-ATPase, and hydrolase domains is shown in **Fig. 5b**. **Figure 5c** provides a detailed view of the predicted conformation of the GDGxNDxP motif in Morex, highlighting the position at which an indel polymorphism introduces a frameshift in *Hv*HMA4 in the genotypes B1K-17-07, WBDC133, HOR4224, HOR14061, Barke, and HOR2180. This motif is thought to function as a hinge that facilitates the transition between the E1 and E2 states of the ATPase enzyme through changes in Mg²⁺ coordination of the first aspartate (D) of the GDGxNDxP. In the E1 state, Mg²⁺ is coordinated by this D residue within the P domain together with residues from the N domain, whereas in the E2 state the N domain interactions are replaced by coordination with residues in the TGES motif of the A domain [30]. Mutations within the GDGxNDxP motif are known to cause Wilson disease (WD) in humans, a disorder characterised by toxic Cu accumulation resulting from impaired Cu transport by the WD protein, a Cu-transporting ATPase and orthologue of *Hv*HMA4. Disease severity depends on the location of the mutation within the protein, with the most severe phenotypes associated with substitutions affecting the first aspartate (D) and the terminal proline (P) residues of the GDGxNDxP motif. When these mutations were recreated in *Escherichia coli*, functional analyses showed that proteins carrying mutations at the D and P residues retained 0% and 10% of wildtype activity, respectively, indicating that both residues are critical for transporter function [30]. Given that the frameshift present within *Hv*HMA4 alleles occurs within the highly conserved GDGxNDxP motif, immediately between residues known to be critical for ATPase function and predicted to disrupt downstream structure, protein function is likely to be severely compromised in genotypes carrying the frameshift allele. Overall, frameshift alleles within this gene family are not uncommon among the surveyed genotypes, and are also present in elite germplasm, where they are likely to result in complete loss of gene function.

### Transcript abundance of *HMA* gene family members in cv. Morex

Numerous barley transcriptomic studies in the public domain have recently been integrated into a web-accessible ‘on demand’ gene expression database (MorexGeneAtlas, https://ics.hutton.ac.uk/morexgeneatlas/index.html) that allows display of gene-specific transcript abundance data (i.e. gene expression) quantified as TPM against the Morex Reference Transcript Dataset (HvMxRTD [25]). Here we focused specifically on five of the component project datasets; a tissue specific developmental series (PRJEB14349), a dissected ‘root zone’ dataset (PRJNA589222), a graduated salt stress experiment (PRJNA639318), seedling tissues exposed to heavy metal stress (PRJNA382490) and in shoots in response to spot botch infection (PRJNA315041). We considered these potentially informative from the perspective of the *HMA*s.

### *HvHMA* expression during development

In order to understand where and when *Hv*HMA proteins were playing functional roles, we explored the expression of *HvHMA* gene family members across 16 dissected tissues representing different stages of barley plant development (PRJEB14349 [31]). Many of the *HvHMA* genes exhibited tissue and temporal-specific expression hinting at diverse and varied physiological roles beyond simple nutrient uptake in the root, as might be expected for members of this gene family (**Fig. 6a**, with TPM values for each *HvHMA* transcript presented in **Supplementary Table S19**). For example, relative to their expression in other tissues *HvHMA9* and *HvHMA12* are highly and specifically expressed in internode and senescing leaf tissues, with expression levels of 102.01 TPM and 33.32 TPM respectively. The tissue-specific elevated expression of these genes suggests specialised roles for *Hv*HMA9 and *Hv*HMA12 proteins in these tissues. Other family members share similar expression patterns. *HvHMA1*, *HvHMA8*, and *HvHMA12* are leaf-specific, with their highest expression levels observed in leaf, senescing leaf, epidermis, and etiolated leaf samples, suggesting potential roles in supplying metal ions to proteins associated with photosynthetic function — roles already established for Arabidopsis orthologues *At*HMA1 and *At*HMA8 [32]. *HvHMA5* and *HvHMA2a* exhibit highest expression in the 28-day-old root (13.14 and 19.57 TPM) and are expressed at their lowest levels in etiolated leaf and inflorescence tissues. Generally, *HvHMA3* had the lowest level of expression (expression ranging from 0.14 TPM in the caryopsis 15 DPA to 5.71 TPM in the senescing leaf) and *HvHMA11* had the highest level of expression (ranging from 10.40 TPM in the caryopsis 15 DPA to 121.21 TPM in the caryopsis 5 DPA) across the tissues and timepoints included here.

**Figure 6:**
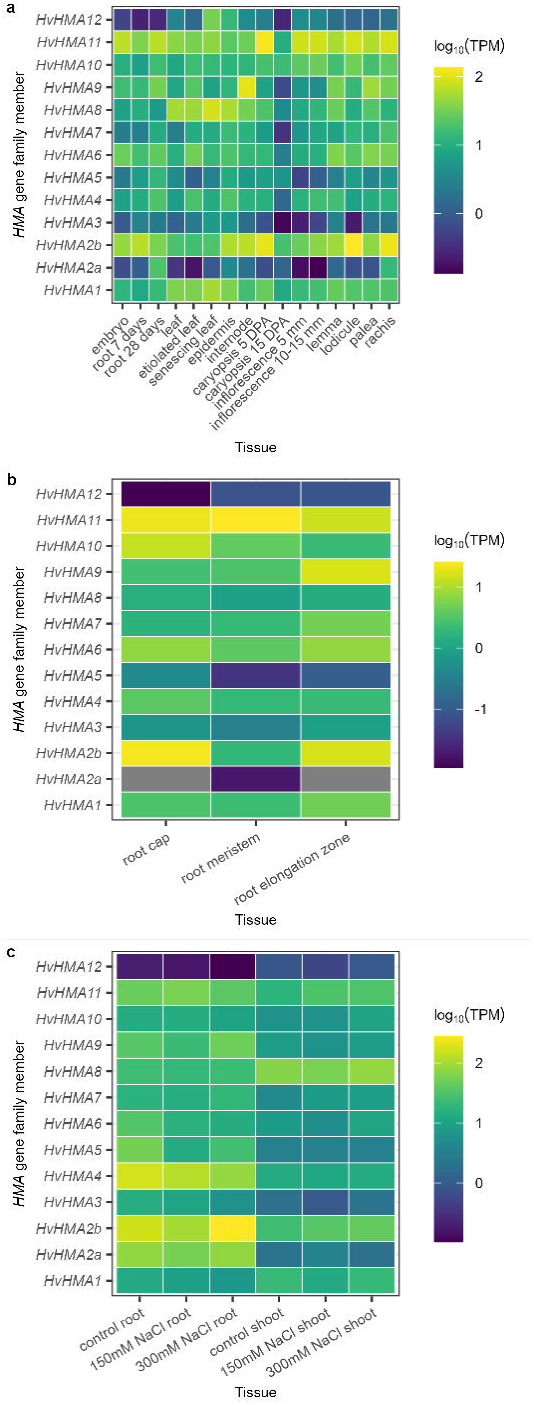
Expression profiles of *HvHMA* family members in cv. Morex. Expression values are shown as logLL-transformed averaged expression levels (in TPM) across multiple biological replicates, with colour indicating relative transcript abundance: yellow represents higher expression, turquoise indicates moderate expression, and dark purple denotes lower expression. Grey indicates the number of reads was below the threshold for quantification. **(a)** Expression profile of the *HvHMA* gene family across 16 developmental stages (*n* = 6 for all tissues except internode where *n* = 7, study accession number PRJEB14349). **(b)** Expression profile of the *HvHMA* gene family in root and shoot tissues under NaCl stress conditions (*n* = 3, study accession number PRJNA639318). **(c)** Expression profile of the *HvHMA* gene family in the root cap, meristem, and elongation zone in three-day-old seedlings (*n* = 3, study accession number PRJNA589222).

### *HvHMA e*xpression across the root tip

Given the general observation of increased expression in the root for most *HvHMAs* we next explored the spatial variation in transcript abundance across three root zones (root cap, meristem, and elongation zone). We observed distinct spatial expression patterns (**Fig. 6b** and **Supplementary Table S20**). *HvHMA11* exhibited the highest overall expression in the root, particularly in the root meristem where it was higher than all other surveyed genes (25.80 TPM). *HvHMA11* was also expressed at a relatively high level in both the root cap (21.57 TPM) and elongation zone (13.94 TPM). *HvHMA2b* was expressed most highly in the root cap (23.15 TPM) and elongation zone (15.97 TPM) but was expressed at low levels in the root meristem (1.93 TPM). *HvHMA9* and *HvHMA10* exhibited relatively high levels of root zone-specific expression, with the former being specifically expressed in the elongation zone (16.88 TPM) and the latter in the root cap (13.13 TPM). *HvHMA3*, *HvHMA5*, *HvHMA8*, and *HvHMA12* showed uniformly low expression levels across the three root tissues. *HvHMA2a* expression could not be detected in the root cap or elongation zone and was negligible in the meristem (0.02 TPM). Analysis of gene expression across the root tip demonstrates that *HvHMA* expression is not only variable between tissues and timepoints, but that individual family members exhibit complex patterns at the sub-tissue level.

### *HvHMA e*xpression under NaCl stress

Sodium is a naturally abundant macronutrient that through a series of complex interactions, including competition for the same uptake channels or the induction of osmotic stress leading to secondary effects on growth, can have widespread impacts on heavy metal homeostasis. We were therefore interested to assess whether Na stress induced a response by any *HvHMA* gene family members. We explored changes in gene expression in the roots and shoots of all 13 *HvHMAs* under control and increasing NaCl stress conditions. In agreement with the developmental expression dataset, all *HvHMA*s but *HvHMA1*, *HvHMA8*, and *HvHMA12* were expressed at higher levels in the root tissues than the shoot tissues under control conditions (**Fig. 6c** and **Supplementary Table S21**). The expression of *HvHMA2a*, *HvHMA4*, and *HvHMA5* was highest in the root tissue under control conditions, while *HvHMA2b* and *HvHMA9* were most highly expressed in root under 300 mM NaCl treatment. *HvHMA2b* and *HvHMA11* displayed a progressive increase in shoot expression across control, 150 mM, and 300 mM NaCl treatments (25.11, 35.95, & 42.89 TPM for *HMA2b*, and 17.65, 30.82, & 31.45 TPM for *HvHMA11*). A pattern of increased expression in response to increased NaCl was not universal; *HvHMA8*, *HvHMA6*, and *HvHMA10* exhibited increased expression in shoot tissues between control and 300 mM NaCl but decreased expression between the control and 150 mM NaCl (**Supplementary Table S21**). In root tissue a similar pattern of increased expression under 300 mM NaCl but decreased expression under 150 mM NaCl was observed for *HvHMA2b*, *HvHMA9*, and *HvHMA7* (**Supplementary Table S21**). Consistent with the low expression pattern observed across the 16 tissues during development (**Fig. 6C**), *HvHMA12* maintained very low expression levels across all conditions, from 0.10 TPM (root, 300 mM NaCl) to 0.92 TPM (shoot, 300 mM NaCl). We conclude that the salt stress-responsive expression of certain *HvHMA* family members, particularly *HvHMA2b* and *HvHMA8* in root and shoot tissues respectively, points to possible roles in mediating the response to NaCl stress.

### *HvHMA* expression in response to heavy metal stress

In response to heavy metal stress, plants dynamically regulate the abundance and localisation of genes which ameliorate the stress condition (for example through metal re-localisation, sequestration, or by producing a mitigating or protective product) which can be reflected in changes in transcript abundance and provide insight into gene function. We therefore examined *HvHMA* transcript abundance in response to heavy metal stress. We observed significant differences in *HvHMA* gene expression response in comparison to control conditions in both seedling root and leaf tissue when plants were exposed to different heavy metal stresses. Several *HMA* genes were differentially expressed between control and zinc (570 μM), copper (50 μM), or cadmium (80 μM) treated root and shoot tissues from 7-day-old seedlings (**Fig. 7**, **Supplementary Table S22**). In roots, *HvHMA2b* and *HvHMA4* were both significantly up-regulated under cadmium stress (logLLFC = 0.83 and 0.93, respectively, *p*-value < 0.05) (**Fig. 7c**). *HvHMA2b* was also significantly upregulated under copper stress (logLLFC = 1.33) as was *HvHMA4* under zinc stress (logLLFC = 0.74). *HvHMA5* showed differential regulation depending on the applied heavy metal: on cadmium treatment it was downregulated (logLLFC = -0.77), but on copper treatment it was up-regulated (logLLFC = 0.64). *HvHMA6* and *HvHMA9* were both down-regulated in roots under zinc stress (**Fig. 7a**), with significant fold-changes of expression (logLLFC = -0.98 and -0.67, respectively). In shoot tissues, *HvHMA11* was up-regulated under zinc and cadmium treatments (logLLFC = 1.06 and 0.84, respectively, *p*-value < 0.05) (**Fig. 7d** and **f**). In the shoot, *HvHMA5* was significantly down-regulated in multiple treatments (logLLFC = -0.90 and -0.96 under cadmium and zinc treatments, respectively, *p*-value < 0.05). *HvHMA1*, *HvHMA9*, *HvHMA8*, and *HvHMA12* were down-regulated under zinc and cadmium stresses in the shoot (**Fig. 7d** and **f**), with *HvHMA12* exhibiting a relatively large decrease in expression level under cadmium treatment (logLLFC = -6.65 and *p*-value = 0.023). Copper stress elicited no significant change in *HMA* gene family expression in the shoot (**Fig. 7e**, **Supplementary Table S22**). Analysis of *HvHMA* transcriptional responses to heavy metal stress support the idea that most family members participate in heavy metal stress responses, as evidenced by significant stress-induced changes in expression in roots and/or shoots. However, this does not definitively exclude a role in heavy metal stress responses for *HvHMA3*, *HvHMA7*, or *HvHMA10*, which do not show such changes; rather, their regulation may occur predominantly at a post-transcriptional level.

**Figure 7:**
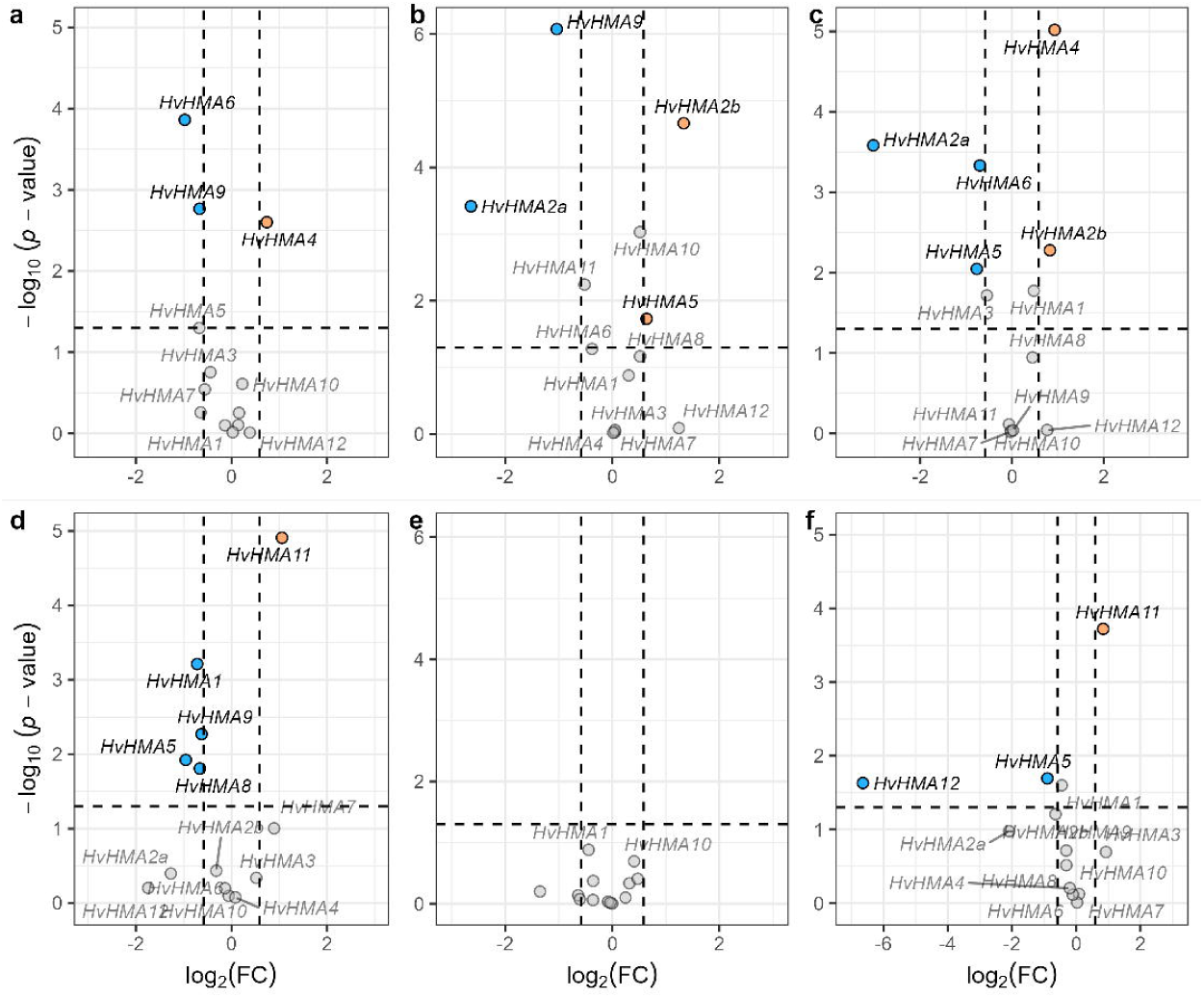
Volcano plots showing differential expression of *HvHMA* gene family members in seven-day-old barley seedlings (*cv.* Morex) exposed to heavy metal-induced stress. (a-c) Root and shoot (d-f) tissues of seedlings treated with 570 μM Zn (a and d), 50 μM Cu (b and e), or 80 μM Cd (c and f). Each plot displays the logL fold change (x-axis) versus statistical significance (−logLL adjusted p-value; y-axis). Genes significantly upregulated (logLFC ≥ 0.585; adjusted p < 0.05) are shown in red, and significantly downregulated (logLFC ≤ −0.585; adjusted p < 0.05) in blue. Expression levels were derived from re-analysis of PRJNA382490.

### *HvHMA* expression in response to spot blotch infection

Plants deploy a number of metal homeostatic mechanisms to control diseases either directly or in association with the general immune signalling responses and we were therefore interested to see if HMA proteins potentially played a role. Members of the *HvHMA* gene family were significantly differentially regulated in shoots in response to infection with *Bipolaris sorokiniana* over time (**Supplementary Fig. S3** and **Supplementary Table S23**). *HvHMA11* was significantly up-regulated (logLLFC = -2.14, 0.97, and 1.70 at 12, 24, and 36-hour post-infection respectively, *p*-value < 0.05) and *HvHMA2a* significantly downregulated between control and infected samples at all timepoints (logLLFC = -3.44, - 2.51, and -1.67 at 12, 24, and 36-hour timepoints respectively, *p*-value < 0.05). Some *HMA* genes exhibited temporal differential expression; *HvHMA9* was strongly up-regulated in infected tissue 24 hours post infection (logLLFC = 2.82, *p*-value = 6.34 x 10^-5^). *HvHMA4* and *HvHMA7* were significantly up-regulated in infected tissues after both 24- and 36-hours post infection (logLLFC = 0.82 & 0.84, and 0.78 & 1.20, for *HvHMA4* and *HvHMA7* respectively at 24- and 36-hour timepoints, *p*-value < 0.05). The expression levels of *HvHMA1*, *HvHMA3*, *HvHMA6*, and *HvHMA10* in the control and infected samples did not change significantly at any timepoint. Our data demonstrate that several *HvHMA*s are dynamically regulated on infection with spot blotch and support a role for *HvHMA*-mediated metal ion transport at the host-pathogen interface.

### *HvHMA* expression level varies significantly by gene identity and genotype across tissues

Next we explored the expression of the *HvHMA* genes in the genotypes of the barley pan-transcriptome dataset (PRJEB64639 [25]) to determine expression level variation between genotypes. The 20 pan-transcriptome genotypes represent germplasm from diverse geographic origins, row types, growth habits, and domestication status (**Supplementary Table S4**) with transcript abundance data collected from five tissues (caryopsis, inflorescence, root, shoot, and coleoptile) as described by Guo *et al.* (2025). Two-way ANOVA revealed significant effects of both genotype and gene identity on *HvHMA* expression level across tissues (**Supplementary Table S24**); however, to be consistent with previous analyses showing predominant root expression, we focused here on the root tissue. As shown in **Figure 8**, substantial variation in root expression level was observed among genotypes, and the degree of variation was not necessarily related to the overall level of expression. For example, *HvHMA2a* exhibited a 39-fold difference in expression between the highest- (ZDM02064, 24.44 ± 7.75 TPM) and lowest-expressing genotypes (HOR3081, 0.62 ± 0.19 TPM), despite being, on average, the second-lowest expressed member of the family. This contrasts with several more highly expressed *HvHMA* genes, for example *HvHMA2b*, which displayed comparatively lower levels of expression variation across genotypes (2.40-fold variation between the lowest and highest-expressing genotypes). In the root tissues, *HvHMA6* exhibited both the highest average level of expression across the genotypes (82.05 ± 24.95 TPM), and the largest range in expression (86.01 TPM). Within the pangenome dataset, OUN333 and Barke were previously identified as carrying frameshift alleles of *HvHMA2a* and *HvHMA4*, respectively. Transcripts of these alleles were expressed at levels comparable to the mean root expression of these genes, with *HvHMA2a* expressed in OUN333 at 2.23 ± 1.01 TPM and *HvHMA4* expressed in Barke at 30.32 ± 3.15 TPM.

**Figure 8:**
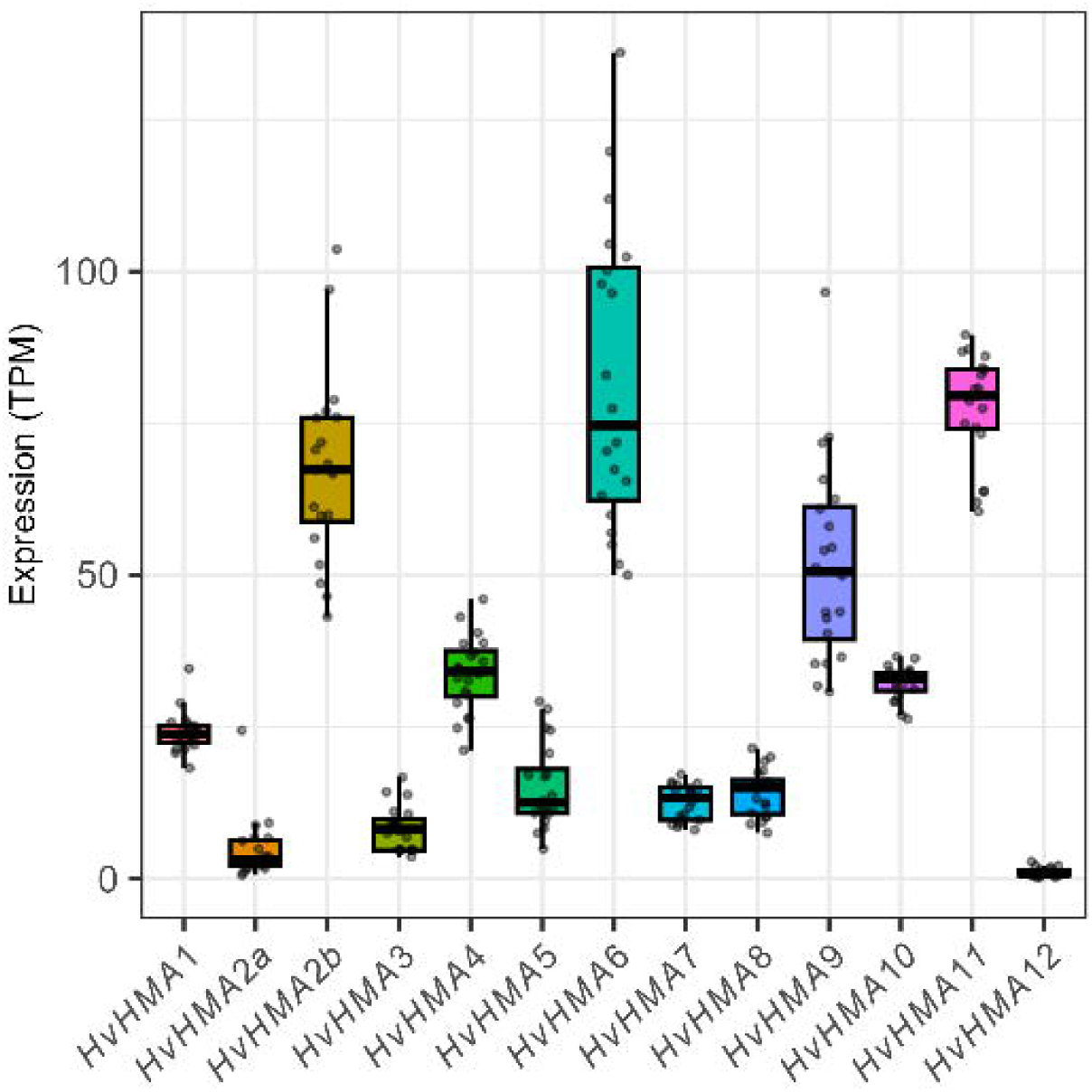
*HvHMA* gene expression in the root tissue of the PanBArt20 lines. Individual grey points represent the average expression value for each genotype (*n* = 3) (Akashinriki, Barke, ZDM02064, ZDM01467, B1K-04-12, Golden Promise, Hockett, HOR10350, HOR13821, HOR13942, HOR21599, HOR3081, HOR3365, HOR7552, HOR8148, HOR9043, Igri, Morex, OUN333 and RGT Planet). Boxplots show the median (centre line), interquartile range (box), and data range excluding outliers (whiskers).

Building on previous allelic analyses, we also examined expression level by allele in the root tissue. However, partly due to the limited number of genotypes represented in the Pantranscriptome dataset, we detected only one significant allele-specific expression level difference, in *HvHMA8*. As shown in **Supplementary Fig. S4**, allele A—present in genotypes such as Barke, Morex, and RGT Planet—exhibited significantly higher expression than allele B, which is found in genotypes including 10TJ18, Akashinriki, and ChikurinIbaraki.

## Discussion

*HMA* gene family size varies considerably depending on identification criteria. In this study, 13 *HMA* genes were identified in barley, distributed across all chromosomes except 1H and 3H. This represents a reduction from the 21 *HMA* genes identified by Zhang *et al.*, (2021), attributable to the implementation of more stringent identification criteria in this study. Specifically, family membership required the presence of both conserved HMA protein domains (E1-E2 ATPase and hydrolase domains) and the defining P_1B_-type ATPase motifs characteristic of the *HMA* subfamily [23,33]. Because the sequences responsible for E1/E2 ATPase and ATP hydrolase activity are highly conserved across species, they provide reliable targets for identifying *HMA* genes. Therefore, a domain-based search approach was used as unlike homology-based methods this approach identifies genes based on functional domain conservation, enabling the detection of more divergent family members that may share low overall sequence similarity but retain core functional motifs. The number of *HMA* gene family members reported here is comparable to that identified in other diploid Poaceae species (nine *HMA*s in *Oryza sativa* and 11 in *Zea mays* [17]) and benefits from using the most current barley genome assembly [24], which provided more complete gene sequences than previously.

*HvHMA2a* and *HvHMA2b* were identified as likely tandem duplicated genes due to their proximity to one another and sequence similarity. Although tandem duplication has been reported within the *HMA* gene family in plants, including in soy bean (*GmHMA1* and *GmHMA2* [21]) and potato (*StHMA19* and *StHMA20* [34]), this is the first report of this evolutionary mechanism in barley. Notably, *HvHMA2a* and *HvHMA2b* exhibit considerable sequence similarity both between themselves and with *AtHMA4*, suggesting they may represent duplicated orthologous genes. This is particularly significant because *AtHMA4* undergoes tandem triplication and quadruplication in the metal hyperaccumulator species *Arabidopsis halleri* and *Noccaea caerulescens*, where the duplications confer hypertolerance to Cd and Zn [35,36]. The maintenance of multiple copies of this sequence within a genome may represent convergent evolution for optimising metal homeostasis. However, the distinct expression profiles of *HvHMA2a* and *HvHMA2b* during development and within the root tip suggest different roles for these genes. When orthologues of these genes were identified in the barley pangenome accessions, it was notable that *HvHMA2a* could be identified in all 76 lines, while *HvHMA2b* was absent in one accession, WBDC133. WBDC133 is a wild barley, and it is therefore possible that WBDC133 diverged from other barleys prior to the duplication event, or that *HvHMA2b* differs greatly at the sequence level in WBDC133. Interestingly, protein structure prediction showed that *Hv*HMA2a and *Hv*HMA2b have large cytosolic regions with low conservation with other known proteins. Further investigation is required to determine the functional relevance of these regions.

Similar to previous studies, phylogenetic analysis divided the *HvHMA* sequences into five groups [23]. However, this analysis also revealed a notable divergence of several *HMA* homologues located on chromosome 4H forming a distinct clade, containing *HvHMA7*, *HvHMA10*, and *HvHMA11*, with weak orthology to *HMA* genes identified in other plant species. This may suggest lineage-specific expansion or accelerated evolution within the barley genome on 4H, potentially arising from ancient segmental duplications unique to Hordeum. Although functional diversification through gene duplication has been reported in Hordeum previously (for example, the *Vrs1*/*HvHox2* [37] and *HvPUB15*/*HvARM1* [38] gene pairs), entirely species-specific *HMA* clades are rarely observed.

The order of domains and motifs in *HMA* proteins is conserved across Poaceae species [17,19]. This arrangement of domains (between zero and three HMA domains, followed by an E1-E2 ATPase domain, and a hydrolase domain) was broadly consistent among the barley *HMA* gene family. Four *HMA* family members in this study contained three adjacent copies of the HMA domain, representing an increase compared to previous barley analyses that reported only *HvHMA5* as having a triplicate HMA domain arrangement [23]. *HvHMA4*-*6*, and *HvHMA9*, which share this configuration, showed strong transcriptional responses to most heavy metal stresses in either root or shoot tissues, suggesting a functional link between this domain architecture and stress regulation. All barley *HMA* genes with three HMA domains were assigned to group A of the *HMA* phylogenetic tree, implying that this structural feature may underlie the evolutionary divergence of this subgroup. Moreover, the triplicate N-terminal HMA domain arrangement is conserved across multiple species, including *At*HMA5, *Os*HMA6 [23], *Zm*HMA5a [19], and *Ta*HMA4 [17], pointing to an ancient evolutionary origin.

The reduction in *HMA* allele diversity from wildtype through landrace to cultivated accessions aligns with the documented allelic bottleneck affecting other barley genes [39]. Several *HvHMA* genes, including *HvHMA1*, *HvHMA2b*, and *HvHMA4*, contained multiple alleles with frameshift mutations. Since these mutations typically render gene products non-functional their presence may indicate that there is some level of functional redundancy between the *HvHMA* genes or between these genes and other loci preventing the loss of accessions with frameshifts alleles. The occurrence of a frameshift allele in elite cultivar Barke (in addition to multiple other genotypes) further supports this hypothesis, as this accession must be able to maintain agronomic performance despite harbouring a probable non-functioning allele. In rice, *Os*HMA4 (sharing 87.97% similarity with *Hv*HMA4) sequesters Cu into root-localised vacuoles; consequently, loss of function leads to elevated Cu levels in aerial tissues, including the grain [40]. However, we propose that *Hv*HMA4 may preferentially transport Zn rather than Cu — consistent with the role of its HMA4 orthologue in Arabidopsis [9,35,41]. This hypothesis is supported by both the significant up-regulation of *Hv*HMA4 in roots treated with Zn (**Figure 7a**) in addition to the analysis of an available grain element profile dataset (previously reported in Houston *et al.*, 2020), in which Barke ranked among the genotypes with the lowest grain Zn concentrations, while its grain Cu concentration was close to the dataset average (**Supplementary Figure S5**).

Here the expression patterns of the *HvHMA* gene family were examined across multiple transcriptomic datasets to elucidate their potential roles in barley. While it might be expected that the transcript abundance of heavy metal transporters would be most dynamic under metal stress or restricted to root tissues (that typically interface with metal gradients), the expression patterns presented here suggest otherwise. Various barley *HMA* genes were significantly differentially expressed under all surveyed conditions, treatments, and developmental stages, indicating that their regulation and roles are far more complex and widespread than previously assumed. This broad transcriptional responsiveness implies multifunctionality, potentially linking *HvHMA*s not only to metal homeostasis but also to broader physiological and developmental processes such as grain maturation and leaf senescence.

When *HMA* expression was surveyed across developing tissues, we observed that *HvHMA11* and *HvHMA2b* were strongly induced in early developing grain, and other reproductive and floral organs. Between the developmental stages of 5 DPA and 15 DPA caryopsis most *HMA* genes were down-regulated, some experiencing a 14-fold reduction in expression (*HvHMA7*). Detterbeck *et al.* (2016, 2020) showed that distinct spatial gradients of metal micronutrients are established during barley grain development. Given that *HMA* family members are known metal transporters in other plant species, it is plausible that these genes are strongly differentially regulated during barley grain development. Several genes, including *HvHMA8*, *HvHMA9*, and *HvHMA12* were most highly expressed in vegetative organs, potentially linking *HMA*s not only to metal detoxification but to broader physiological and developmental processes. In Arabidopsis, members of the *HMA* gene family have been linked to ethylene signalling (*AtHMA7*) and photosynthesis (*AtHMA6*), through their capacity to deliver metal ions which act as key co-factors to enzymes and receptors in these pathways [43,44].

Adding to the spatial complexity of *HMA* expression in barley, transcriptomic analysis within the barley root revealed that *HMA* gene expression is regulated within discrete tissue zones.

*HvHMA9* showed elevated transcript abundance specifically in the root elongation zone, with minimal expression detected in the root meristem or cap. In rice roots, investigations of *OsHMA2*, *OsHMA4*, and *OsHMA5* have revealed high expression levels specifically confined to root pericycle cells [13,40,45]. Due to how roots were partitioned for gene expression analysis in the dataset here, it is impossible to assess whether barley orthologues of these genes are specifically expressed in root pericycle cells. However, *HvHMA2b*, *HvHMA4*, and *HvHMA5* were found to be most highly expressed in the root cap tissue. The specificity of HMA expression to discrete root zones is consistent with the root’s role as the primary interface between the plant and metal gradients. Plant metal transporters have been shown to exhibit highly specific spatial localisation, down to the level of cellular polarisation, as demonstrated for IRON-REGULATED TRANSPORTER 1 (IRT1) in Arabidopsis, which localises to the outer plasma membrane of root epidermal cells to facilitate iron uptake [46].

Additional insight into the role of *HMA* gene family in barley was gained through the analysis of expression data under NaCl and heavy metal stress. Salt and heavy metal stress responses often share regulatory pathways in plants and can occur concomitantly (for example in arid, costal, or polluted areas), making the analysis of these stresses more relevant in conjunction with each other [47–49]. Both stress types can disrupt ion homeostasis and generate reactive oxygen species, requiring common detoxification mechanisms and causing similar symptoms.

Some parallels exist between the expression data of *HvHMA*s presented here under these abiotic stresses. Apart from *HvHMA8* and *HvHMA12*, the *HMA* gene family appeared to be expressed at higher levels in the root tissues than the shoot tissues regardless of NaCl stress levels. Under heavy metal treatments, more genes were significantly differentially regulated in the root tissues than the shoot tissues. Both observations indicate that this family is potentially more active in the root tissues at the interface with the abiotic stress than in aboveground tissues.

The transcript abundance levels of *HvHMA1*-*5*, *HvHMA9*, and *HvHMA11* are both dynamic in NaCl stressed tissues and significantly different under various heavy metal stresses. However, their pattern of expression was not necessarily consistent under both stress conditions. It was found that *HvHMA4* was significantly up-regulated in the root under Cd and Zn stress (in agreement with *OsHMA4* expression patterns [40]), yet in the same tissue decreased transcript abundance was observed with increasing NaCl stress. This broad transcriptional responsiveness implies multifunctionality of these genes and would benefit from transcriptional studies of the *HMA* family in plants exposed to both stresses simultaneously, similar to a study of *Mesembryanthemum crystallinum* [48].

Zhang *et al.* (2021) examined the relative expression of several *HvHMA* family members in response to Cd stress (50 and 100 µM) and observed a stepwise increase in the expression of *HvHMA2b* (denoted by Zhang *et al.* (2021) as *HvHMA1*) and *HvHMA3* in seedling leaves with increasing external Cd concentrations. *HvHMA9* (denoted by Zhang *et al.* (2021) as *HvHMA4*) expression was also elevated under 100 µmol L⁻¹ Cd treatment compared to the control, but not at 50 µM. In contrast, *HvHMA4* (denoted by Zhang *et al.* (2021) as *HvHMA2*) and *HvHMA1* (denoted by Zhang *et al.* (2021) as *HvHMA6*) expression levels decreased under Cd stress. The experimental conditions used by Zhang *et al.* (2021) differ from those used to generate the data analysed in this study in terms of Cd concentration, genotype, and seedling developmental stage.

The importance of nutrient metals in plant pathology has been recognized by farmers since the late 1800s, when copper-based Bordeaux mixture was developed for disease control [50]. Recently, the concept of nutritional immunity has emerged as a discipline focused on how hosts regulate nutrient availability to restrict pathogen growth, either by limiting access to essential nutrients or directing toxic levels of metals towards invading pathogens [51,52]. Although the molecular basis of nutritional immunity in plants is not fully understood, studies show that plants actively alter their ionomes in response to infection, sometimes in a pathogen-specific manner [53–55].

Previous studies indicate that members of the *HMA* gene family are involved in plant pathology. In rice, alleles of *OsHMA5* with decreased root-to-shoot Cu transport capacity have been implicated in increased susceptibility to several viruses [56]. In Arabidopsis, *AtHMA2* and *AtHMA4* are required for resistance to fungal infection, as they mediate the re-localisation of zinc to the infection site [57]. The data analysed here supports an active role for the barley *HMA* gene family in response to infection with spot-blotch, which may be related to this gene family’s role in heavy metal homeostasis. *HvHMA2a*, *HvHMA2b, HvHMA4*-*5*, *HvHMA8*-*9*, *and HvHMA11-12* are all significantly up- or down- regulated under at least one heavy metal stress condition and at least one timepoint after spot blotch infection. Further experiments examining the expression of these genes in barley infected with different pathogens could help identify which are involved in the non-specific immune response, providing targets for developing durable resistance against a wide range of pathogens.

The findings presented here represent the first comparative analysis of HMA gene expression across both tissues and accessions. We observed significant variation in the expression levels of *HMA* family members among the PanBaRT20 barley genotypes, suggesting that the underlying *cis*-regulatory networks differ between them. These findings indicate that breeding strategies could not only target the incorporation of favourable alleles but also harness natural variation in gene expression for trait improvement. This may be particularly valuable for optimising complex traits like metal nutrient homeostasis, where subtle adjustments in expression could be more effective than drastic genetic changes. Previous studies in barley highlighted that modifying *HvHMA3* expression levels through integrating a naturally-occurring transposable element into the gene’s *cis*-regulatory region significantly reduced grain Cd accumulation, demonstrating that expression-based breeding approaches are viable alternatives to coding sequence alterations [14]. However, some caution is warranted when interpreting genotypic differences in expression, particularly in inflorescence and caryopsis, as tissues were collected at matched developmental stages rather than at fixed time points after planting, meaning collection timing may have varied between genotypes.

## Conclusion

While expression analyses provide insights into *HMA* gene function in barley, definitive functional characterization requires additional experimental approaches. Studies in rice and Arabidopsis have employed gene knockout mutants and heterologous expression systems to determine transport specificity of individual family members [13,58]. Analogous studies in barley would identify genes relevant to key breeding objectives, including tissue metal loading, nutrient uptake efficiency, and heavy metal stress tolerance. Following functional characterization and prioritization of *HMA* genes based on relevance to traits of interest, the sequence and expression level variation identified in pan-transcriptome and pangenome resources could be exploited through targeted crosses to integrate potentially beneficial alleles from diverse germplasm into elite cultivars.

## Methods

### Identification of *HMA* gene family members in barley

To determine barley sequences which belonged to the *HMA* gene family, the HMMER profiles related to the conserved domains of *HMA* proteins (E1–E2 ATPase: PF00122; Hydrolase: PF00702) were downloaded from the Pfam database [59] and used to screen the amino acid sequences of the Morex V3 genome assembly (both high and low confidence sequences) (Hv_Morex.pgsb.Jul2020.aa.fa) accessed from the e!DAL - Plant Genomics & Phenomics Research Data Repository [24]. E1–E2 ATPase and Hydrolase domains were considered present in an amino acid sequence if the ‘full sequence’ had an E-value below 0.00001. Amino acid sequences with these domains were then downloaded from EnsemblPlants using the BioMart tool (42 genes, 46 different amino acids). The DKTGT[LIVM][TIS], GDGxNDxP, PxxK, and S/TGE motifs are found in all P-type ATPases (including P_1B_-ATPases, to which *HMA*s belong) and therefore the amino acid sequences were further refined to 34 loci (37 amino acids) that contained these motifs. P_1B_-ATPases are defined from other P-type ATPases by the presence of a CPx (or SPC) motif [33] and a HP motif [8,21]. The list of genes was further refined to those which contain these motifs in addition to the P1B-type ATPase-associated motifs (13 loci, 15 amino acids). **Supplementary Table S1** summarises the molecular characteristics of the canonical transcripts of these 13 loci, including predicted isoelectric point and molecular weight, and lists the genomic locations of the corresponding coding genes. Identification of all motifs was done manually using Geneious Prime® 2022.2.2 (https://www.geneious.com). A reciprocal BLASTP (v. 2.9.0+) [60] of the identified barley proteins against Arabidopsis proteins was conducted using the TAIR database with default parameters to support their assignment to the HMA family. The highest-scoring Arabidopsis alignment for each analysed barley protein, based on bit score and E-value, was annotated as “Heavy Metal Associated” or “Heavy Metal ATPase,” except for HORVU.MOREX.r3.4HG0419450.1 and HORVU.MOREX.r3.4HG0339640.1, which were annotated as H(+)-ATPase and Ca2+-ATPase, respectively. Domains and motif positions within the *HMA* gene family were visualised using ggplot2.

### Designation of gene *HMA* gene names in barley

A Clustal Omega alignment (v. 1.2.2) of *HMA* sequences from Arabidopsis, Rice, and Wheat was conducted using Geneious Prime® 2022.2.2 (https://www.geneious.com) and the results of this alignment allowed the identified barley *HMA* sequences to be designated gene names *HvHMA1* to *HvHMA12* based on sequence similarity with orthologues in these species.

### Prediction of *HMA* protein attributes in barley

The subcellular localisation of barley HMA proteins was predicted using WoLF PSORT (https://wolfpsort.hgc.jp/; accessed on 10 December 2024). Molecular weight and isoelectric points for *HMA* proteins were estimated using Geneious Prime® 2022.2.2 (https://www.geneious.com).

### *Hv*HMA protein conformation prediction

The *Hv*HMA protein sequences from Morex were input into the AlphaFold server and their structure was predicted using the AlphaFold3 model with the seed set to 2 and all other settings set to default. For each HMA protein the model with the highest confidence was visualised using The PyMOL Molecular Graphics System, Version 3.0 Schrödinger, LLC.

### Construction of a phylogenetic tree of *HMA* gene family members

The HMA protein sequences of barley (cv Morex), *Oryza sativa*, *Triticum aestivum*, *Zea mays*, *Sorghum bicolor*, and *Arabidopsis thaliana* were aligned with the online MAFFT v7 tool (https://mafft.cbrc.jp/alignment/software/) using default settings (including the L-INS-i strategy option). Accession numbers, Uniprot IDs, and references for each *HMA* sequence are provided in **Supplementary Table S25**. The best-fit substitution model was determined to be JTT+I+G4+F using ModelTest-NG v0.1.7 [61] and tree inference was conducted according to this model using RAxML-NG v1.1.0 [62]. Non-parametric bootstrapping was conducted using RAxML-NG and bootstrapping converged after 350 replicates using the default MRE-based bootstrapping test cut-off of <0.03. Felsenstein’s bootstrap proportions were computed as the support metric for phylogenetic tree branches.

### Extraction of *HMA* gene family variants across the Barley Pangenome v.2 and determination of *HMA* expression from publicly available RNA seq datasets (*cv.* Morex) & PanBaRT20

The Barley Pangenome v.2 (BPGv2 [26]) and Barley Pantranscriptome (PanBart20 [25]) were used to extract variants of the *HMA* gene family from 76 genome assemblies, determine the expression of *HMA* genes in publicly available RNA seq datasets (*cv.* Morex), and determine of the expression of *HMA* genes in PanBaRT20 accessions. The genomic and canonical transcript sequences for each *HMA* gene family member were downloaded from Morex v3.0 using the Ensembl plants genome browser. The genomic sequences were used in a BLASTn search against the BPGv2 genome sequences and filtered for >95% percentage identity and a total percentage coverage of >70%. A second BLASTn search with the transcript sequence against the PanBart20 transcriptome was carried out, filtering for a percentage identity of >95% and a percentage coverage of >60%. The genomic sequences were extracted using genomic coordinates with bedtools getfasta [63], aligned to the Morex input sequence using minimap2 (-x asm20 -N 0, (Li, 2018)) followed by variant calling with the Naïve Variant Caller tool (Blankenberg D, et al. *In preparation* https://github.com/blankenberg/nvc/). The code for this workflow is available at https://github.com/SchreiberM/BarleyPangenomeVariantExtract. Nucleotide sequences were translated into protein sequences using Geneious Prime® 2022.2.2 (https://www.geneious.com) with default parameters. To emphasize functional relevance, in this study unique alleles were defined by differences in predicted protein sequences. Alleles of each *HMA* family member were designated an alphabetical identifier (A through to AF) in order of most to least common within the accessions of the Barely Pangenome. For 20 of the barley lines (PanBaRT20), for which genotype-specific reference transcript datasets were available, the expression of each of the *HvHMA* genes was retrieved in transcripts per million on a per gene basis from five sampled tissues (caryopsis, inflorescence, root, shoot, and coleoptile). Coleoptile tissue was sampled at four to eight days post-germination, shoot and root tissues at 14 to 17 days post-germination, inflorescences at Waddington stages W6 to W7, and caryopses at 15 to 20 days post-anthesis. Data was retrieved directly from the ‘MorexGeneAtlas’ [25] and Raw data for the publicly available RNA-seq studies are available through the ENA European nucleotide archive (study accession numbers PRJEB14349, PRJNA639318, PRJNA589222).

### Analysis of publicly available RNA seq data for determination of differential expression

The RNA-seq data are available at the NCBI BioProject archive under accession numbers PRJNA315041 and PRJNA382490, respectively. Expression profiles in these resources were generated from 14-day-old seedlings for the former dataset and seven-day-old seedlings for the latter. In both cases, the transcript abundances from RNA-seq reads were estimated using Salmon version 1.2.0 [65] with the PanBaRT transcript reference dataset [25] in the mapping-based mode. Compressed quantified gene expression files were imported into 3D-RNA seq web-based application (https://3drnaseq.hutton.ac.uk/) for RNA-seq analysis [66]. Transcripts were considered expressed if at least one sample had an expression level ≥ 1 count per million (CPM). Transcripts below this threshold were excluded from further analyses. Read counts for the remaining transcripts were normalized with weighted trimmed mean of M-values to log_2_-CPM [67]. A gene was considered expressed if at least one of its transcripts met the above expression criterion. Genes were considered differentially expressed if logL(FC) ≥ 0.585 or logL(FC) ≤ –0.585 (≈ 1.5-fold up- or down-regulation) and adjusted *p*-value < 0.05, in line with thresholds which have previously been reported as biologically relevant [68].

## Data Availability Statement

The datasets analysed in the current study are available in the ENA European nucleotide archive (https://www.ebi.ac.uk/ena/browser/) under BioProjects PRJEB14349, PRJNA639318, PRJNA589222, PRJNA315041, and PRJNA382490. Sequence data supporting the pan-genomic dataset analysed in this study can be accessed under BioProjects <u>PRJEB40587</u>, <u>PRJEB57567</u>, and <u>PRJEB58554</u> and was generated by Jayakodi *et al.* 2024. Raw Illumina RNA-seq data and Iso-seq CCS reads associated with the pan-transcriptomic dataset analysed in this study was generated by Guo *et al.* 2025 and can be accessed under BioProjects PRJEB64639 and PRJEB64637. The pan-transcriptomic dataset can also be accessed via EoRNA (https://ics.hutton.ac.uk/panbart20/index.html).

## Acknowledgements

KH, MS, JR, and RW acknowledge the support of the Rural and Environmental Science and Analytical Services division of the Scottish Government. JS would like to acknowledge funding from the Mynlefield Trust.

## Figure legends

**Supplementary Figure S1: Unrooted phylogenetic tree of *HMA* protein sequences from barley (red) identified in this study and previously reported sequences from rice (magenta), wheat (green), sorghum (orange), maize (purple), and Arabidopsis (blue).** The analysis revealed five distinct subgroups (A-E) within the HMA gene family. Numbers at branch nodes represent Felsenstein’s bootstrap proportions (expressed as percentages), which indicate the statistical confidence of each branch.

**Supplementary Figure S2:** Schematic representation of the predicted protein products encoded by frameshift alleles of (A) *Hv*HMA1, (B) *Hv*HMA2a, (C) *Hv*HMA2b, (D) *Hv*HMA7, (E) *Hv*HMA8, and (F) *Hv*HMA11, compared with the corresponding proteins encoded by the allele present in the reference cultivar Morex. Grey bars represent the full-length protein sequence, with coloured boxes indicating conserved domains and motifs. Red regions denote amino acid sequences encoded downstream of the frameshift mutation. The positions of the E1–E2 ATPase domain, hydrolase domain, and conserved HMA motifs are shown according to the legend of Figure 5. Frameshift alleles result in premature truncation and/or disruption of conserved functional domains and motifs to varying extents. Amino acid positions are indicated on the x-axis.

**Supplementary Figure S3: Differential expression of *Hv*HMA genes in spot blotch–infected barley shoots.** Volcano plots show *HvHMA* gene expression in 14-day-old barley (*cv.* Morex) shoots at (a) 12, (b) 24, and (c) 36 hours post-infection compared with mock controls. Each plot displays logL fold change (x-axis) versus statistical significance (−logLL adjusted p-value; y-axis). Genes significantly upregulated (logLFC ≥ 0.585, adjusted p < 0.05) appear in red; significantly downregulated genes (logLFC ≤ −0.585, adjusted p < 0.05) appear in blue. Grey points indicate genes not meeting significance thresholds. Expression values were obtained from publicly available RNA-seq data (*n* = 4; study accession PRJNA315041).

**Supplementary Figure S4: *HvHMA8* expression level by allele in the root tissue of barley pan-transcriptome genotypes.** Individual points represent the average expression value for each genotype (*n* = 3) (Akashinriki, Barke, ZDM02064, ZDM01467, B1K-04-12, Golden Promise, Hockett, HOR10350, HOR13821, HOR13942, HOR21599, HOR3081, HOR3365, HOR7552, HOR8148, HOR9043, Igri, Morex, OUN333 and RGT Planet). Boxplots show the median (centre line), interquartile range (box), and data range excluding outliers (whiskers). Letters above boxplots indicate statistical groupings from Tukey’s HSD post hoc test (adjusted p < 0.05); alleles with different letters are significantly different.

**Supplementary Figure S5:** Boxplots representing the concentrations of (a) Zn and (b) Cu in the grain of a two-row spring barley population (*n* = 130) previously published by Houston *et al.* 2020. Genotypes are ordered by median concentration (in parts per million, ppm) from lowest to highest for each element. For each genotype, element concentration of five biological and five technical replicates was quantified. Box boundaries represent the 25th and 75th percentiles, central lines show medians, and dots represent outliers. The boxplot representing data for genotype Barke is coloured in red.

