## Supplementary figures and images for "Pan-genomic and pan-transcriptomic analysis of the Heavy Metal ATPase family reveals diverse expression patterns and functional roles in barley"

### Figure S2.png

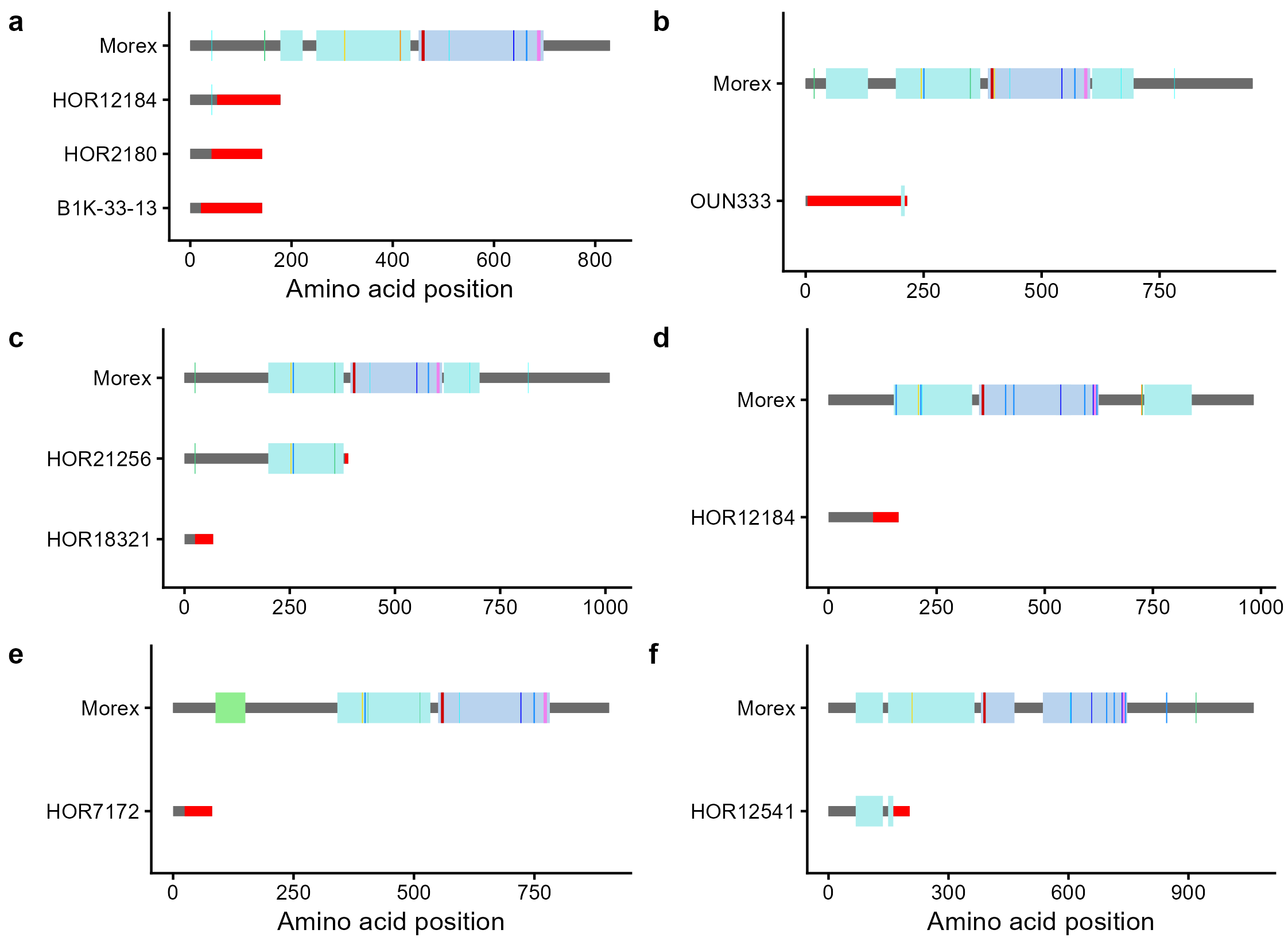

### Figure S3.png

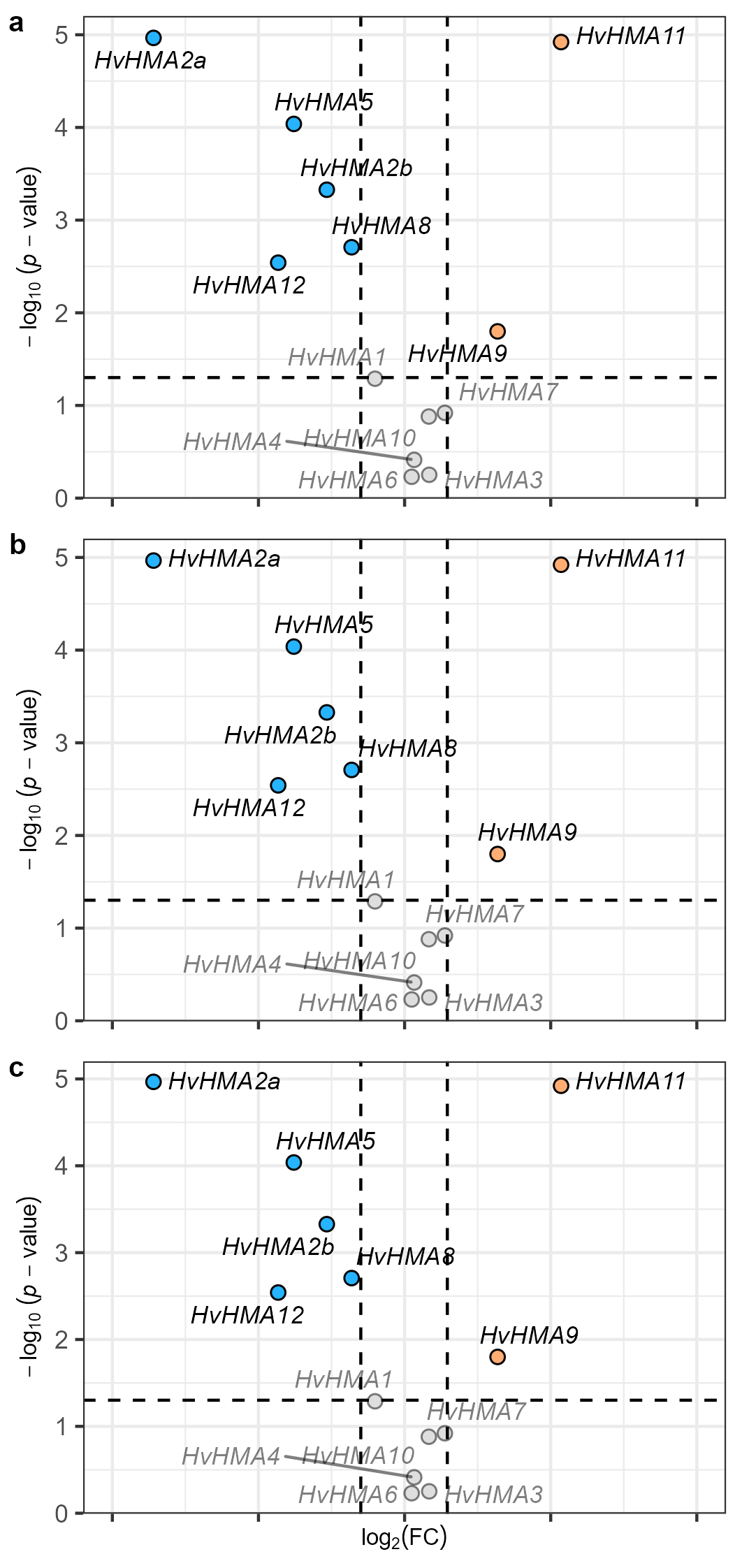

### Figure S4.png

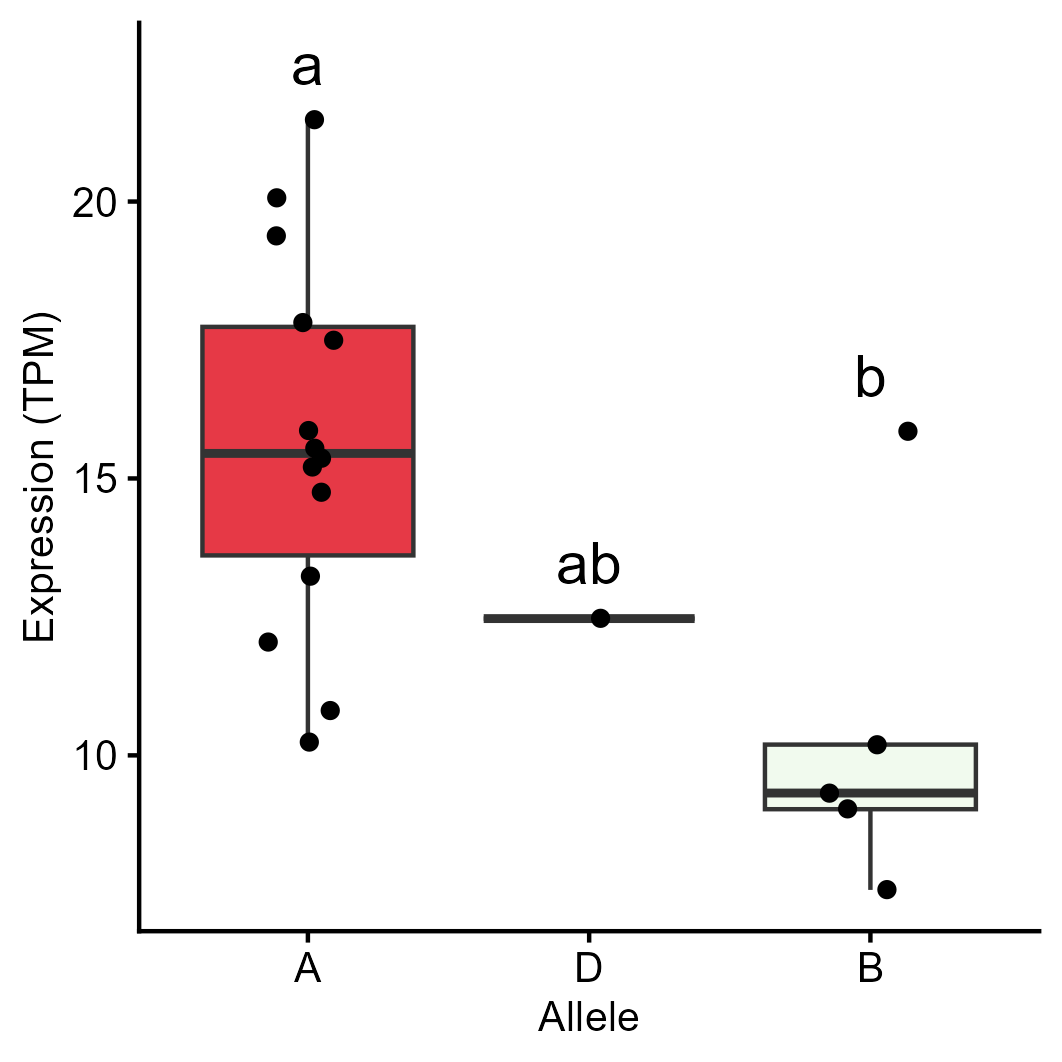

### Figure S5.png

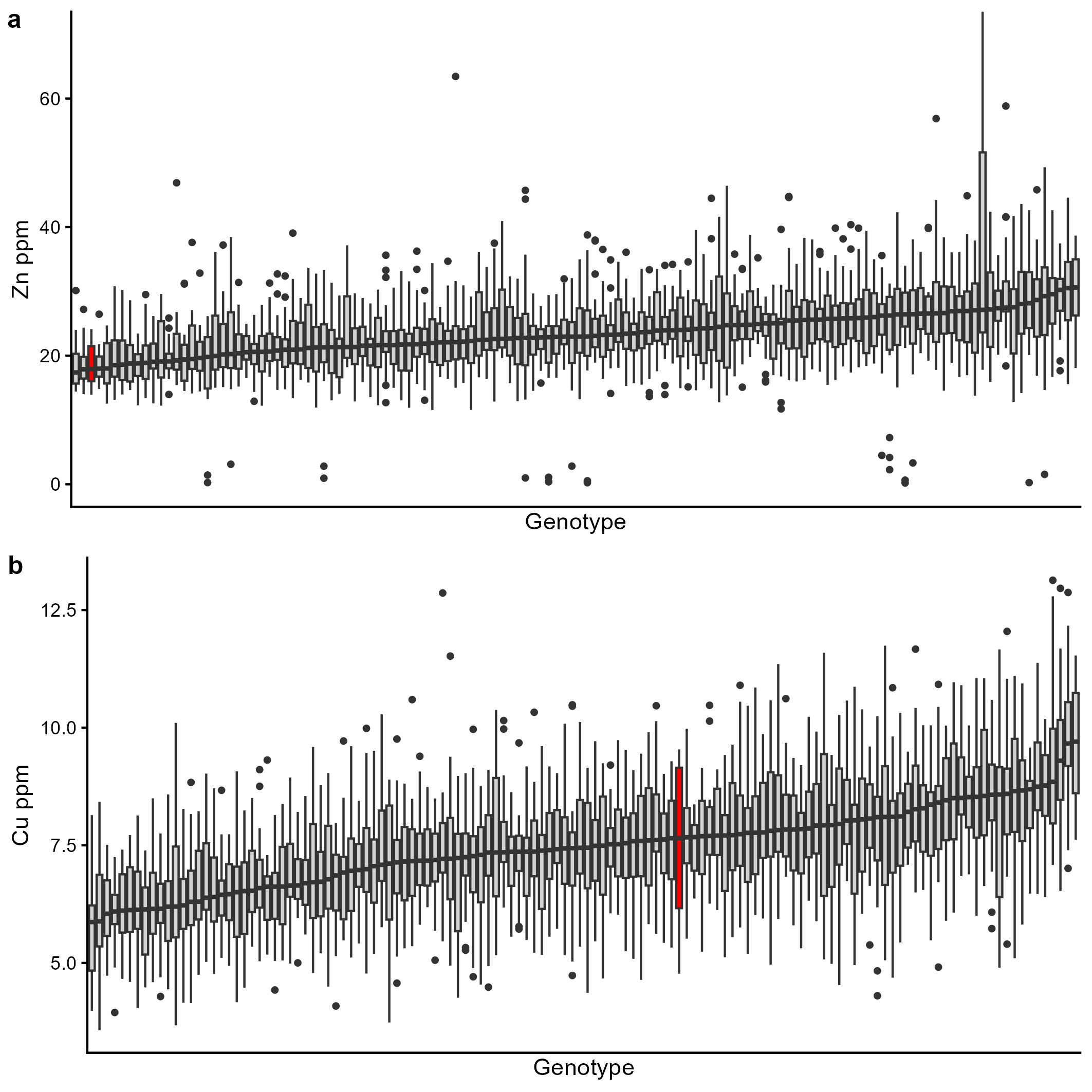

### SFigure 1.jpg

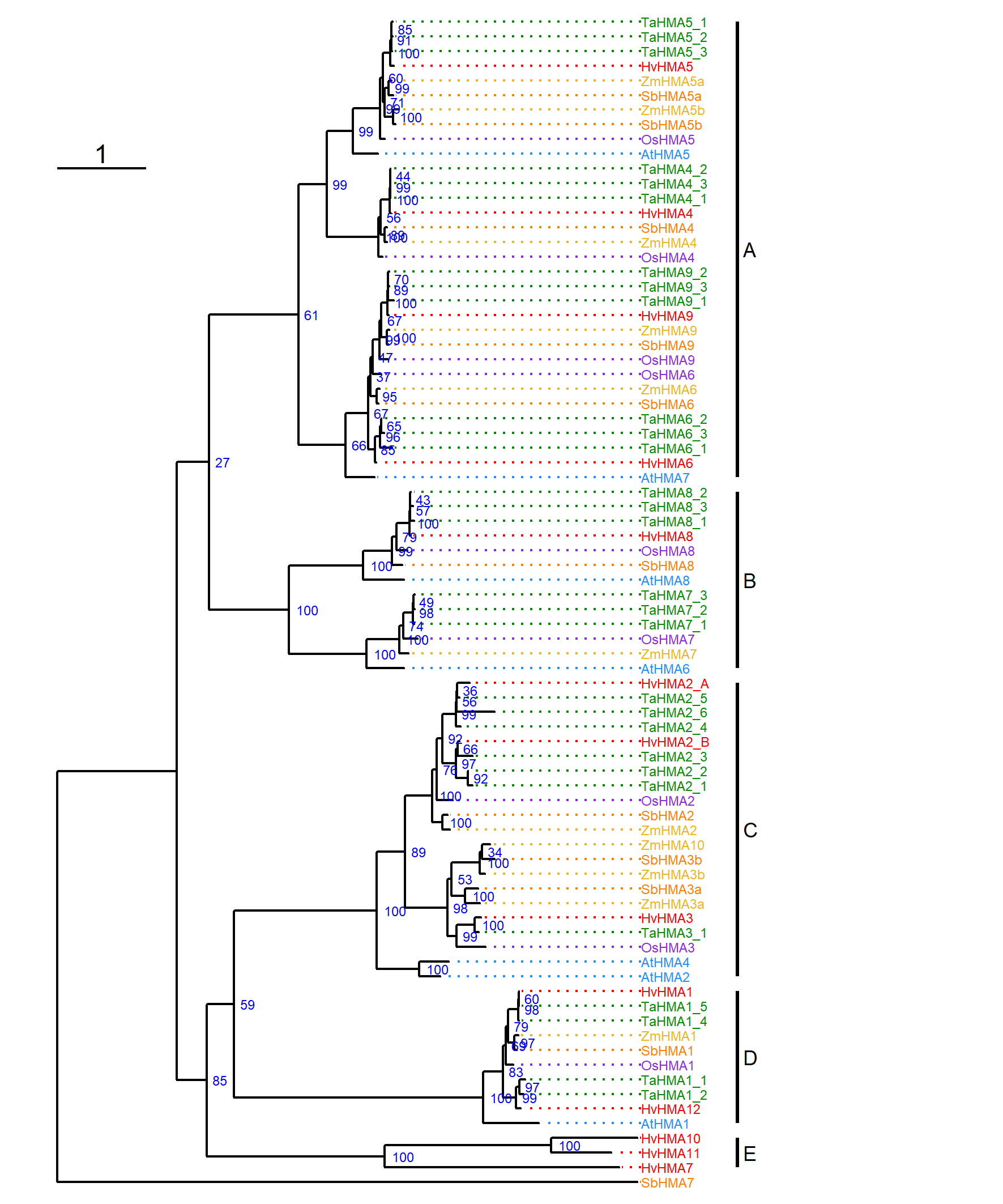
